# Recognition determinants of improved HIV-1 neutralization by a heavy chain matured pediatric antibody

**DOI:** 10.1101/2023.01.19.524673

**Authors:** Sanjeev Kumar, Swarandeep Singh, Arnab Chatterjee, Prashant Bajpai, Shaifali Sharma, Sanket Katpara, Rakesh Lodha, Somnath Dutta, Kalpana Luthra

## Abstract

The structural and characteristic features of HIV-1 broadly neutralizing antibodies (bnAbs) from chronically infected pediatric donors are currently unknown. Herein, we characterized a heavy chain matured HIV-1 bnAb 44m, identified from a pediatric elite-neutralizer. Interestingly, in comparison to its wild-type AIIMS-P01 bnAb, 44m exhibited moderately higher level of somatic hypermutations of 15.2%. The 44m neutralized 79% of HIV-1 heterologous viruses (n=58) tested, with a geometric mean IC_50_ titer of 0.36 µg/ml. The cryo-EM structure of 44m Fab in complex with fully-cleaved glycosylated native-like BG505.SOSIP.664.T332N gp140 envelope trimer at 4.4Å resolution revealed that 44m targets the V3-glycan N332-supersite and GDIR motif to neutralize HIV-1 with improved potency and breadth, plausibly attributed by a matured heavy chain as compared to that of wild-type AIIMS-P01. This study further improves our understanding on pediatric HIV-1 bnAbs and structural basis of broad HIV-1 neutralization by 44m may be useful blueprint for vaccine design in future.

## INTRODUCTION

Extensive efforts are currently ongoing worldwide to develop a safe and effective human immunodeficiency virus-1 (HIV-1) vaccine. During HIV-1 infection, neutralizing antibodies (nAbs) are elicited against the envelope (env) glycoprotein gp160^1–3^. Highly potent broadly neutralizing antibodies (bnAbs) are found to develop and evolve in only top 1% of HIV-1 infected individuals, classified as elite-neutralizers^4, 5^. So far, seven distinct epitopes of HIV-1 bnAbs have been identified to be present on the viral env that are: V2-apex, V3-glycan N332-supersite, CD4 binding site (CD4bs), silent-face center (SFC), membrane-proximal external region (MPER), gp120–gp41 interface and fusion peptide (FP)^3^. The prime goal is to design and develop an HIV-1 vaccine capable of triggering naïve B cells and steer them to evolve into bnAb generating B cells upon immunization/vaccination^6–8^.

HIV-1 infected infants have been shown to develop bnAb responses within one year of age^9, 10^ while in infected adults, it takes at least 2 to 3 years post infection for the development of bnAbs^4, 11, 12^, suggesting distinct maturation pathways of the bnAbs evolving in children^9, 13^. The HIV-1 bnAbs isolated from adults have been extensively characterized, both structurally and functionally; with some of them exhibiting characteristic features of high somatic-hypermutations (SHM) and long CDR3 regions^3^, while there is a paucity of such information on the bnAbs generated by HIV-1 infected children. A number of studies carried out on HIV-1 infected pediatric cohorts have reported HIV-1 plasma bnAb responses targeted at multiple env epitopes including the V2-apex, N332-glycan supersite, CD4bs and MPER^9, 10, 14–21^; however only two pediatric HIV-1 bnAbs have been reported thus far: BF520.1^10^ and AIIMS-P01^22^. Binding of both AIIMS-P01 and BF520.1 are dependent on the N332-supersite epitope present at the base of the V3-glycan region^10, 22^. Both pediatric bnAbs exhibit limited SHM (∼7%); however, there is paucity of information towards understanding whether an increased SHM can be acquired in bnAbs evolving in HIV-1 infected children and further, if the increased SHM in pediatric bnAbs can lead to increase in their potency and breadth of viral neutralization, as observed in bnAbs evolving in HIV-1 infected adults^1, 3, 23–32^.

Immunogenetic information of HIV-1 bnAbs from adults and children derived from deep sequencing or single cell analysis of B cell repertoire (BCR) can further our understanding of their natural development during the course of infection and development of blueprints for rational vaccine design and effective vaccination strategies^6, 7^. Furthermore, structural characterization of potent bnAbs, in complex with native-like trimeric env can provide useful mechanistic insights for broad and potent neutralization of HIV-1 heterologous viruses.

We previously reported the isolation of a bnAb AIIMS-P01, from an antiretroviral naïve HIV-1 clade-C chronically infected pediatric elite-neutralizer AIIMS_330^15, 20, 22, 33, 34^. Recently, for the first time, we identified several adult HIV-1 bnAbs clonotypes targeting multiple epitopes in a pair of monozygotic twin pediatric elite-neutralizers AIIMS_329 and AIIMS_330 from longitudinal (3 time points samples) bulk B cell repertoire analysis by next-generation sequencing (NGS)^35^. Herein, to delineate the characteristics of an affinity matured lineage member antibody of our previously discovered AIIMS-P01 pediatric bnAb, we performed the structural and functional characterization of a heavy chain matured pediatric HIV-1 AIIMS-P01 bnAb lineage monoclonal antibody clone 44m (referred to as 44m). The 44m exhibited moderate level of SHM (15.2%) and demonstrated near about 79% HIV-1 neutralization breadth with ∼2 times improved potency than AIIMS-P01 wild-type (WT) bnAb. Cryo-EM analysis of 44m in complex with native-like fully cleaved and glycosylated BG505.SOSIP.664.T332N gp140 trimer revealed structural insights that may attribute to its neutralization breadth.

## RESULTS

### Identification of a matured AIIMS-P01 lineage antibody

We previously reported the isolation and characterization of a broad and potent anti-HIV-1 bnAb AIIMS-P01 from an Indian clade-C infected pediatric elite-neutralizer AIIMS_330^22^. Recently, for the first time, we identified several adult HIV-1 bnAbs clonotypes targeting multiple epitopes in monozygotic twins pediatric elite-neutralizers AIIMS_329 and AIIMS_330 from longitudinal (3 time points samples) bulk B cell repertoire analysis by NGS^35^. Based on our analysis pipeline we identified few (n=21) matured AIIMS-P01 bnAb heavy chain lineage members/clonotypes (**Figure 1A and S1; Table S1 and S2**) from the year 2018 time point of the AIIMS_330 pediatric elite-neutralizer, from whom AIIMS-P01 was isolated previously^20, 22, 35^. These 21 heavy chain sequences shared the same clonotype with AIIMS-P01 pediatric HIV-1 bnAb, with varied and moderate level (30 nt – 47 nt) of somatic hypermutations (SHM) (**Table S2**). The clonotypes were defined as sequences sharing the same V and J genes, having same CDRH3 length and more than 80% CDRH3 identity (**Table S2**). Like AIIMS-P01 WT bnAb, all 21 matured lineage members maintained the presence of 5 amino acid (AA) indel in the heavy chain framework region 3 (FRH3), however, the mature lineage members exhibit the indel ‘SDPIR’ instead of ‘SNPSR’ which is present in the AIIMS-P01 WT bnAb (**Figure S1 and Table S2**)^22^. Next, we calculated the antibody mutation probabilities using ARMADiLLO, an online server developed to identify both improbable and probable mutations^36^. Interestingly, we found the presence of 4 improbable mutations with less than 2% frequency in the 44m antibody (**Figure S2**) when we analyzed the 44m sequence using ARMADiLLO method^36, 37^. The CDRH3 region of the matured lineage members showed the presence of varied mutations (**Figure 1B and Table S2**). We did not find matched CDRL3 light chain genes as that of the AIIMS-P01 from our sequencing data. As observed by us previously, the heavy chain, specifically heavy chain 5 AA indel dominantly contributed to recognition of N332-supersite, HIV-1 neutralization breadth and autologous virus neutralization^22^. Therefore, to evaluate the effect of higher level of SHM in matured AIIMS-P01 lineage members, herein, we synthesized a matured (15.2% SHM) AIIMS-P01 bnAb heavy chain gene with >80% similar CDRH3 sequence as of AIIMS-P01, designated as 44m and expressed the mAb by co-transfecting plasmids carrying this heavy chain gene and the WT AIIMS-P01 light chain gene. The SHM in 44m antibody heavy chain gene was compared to AIIMS-P01 WT and other HIV-1 bnAbs (**Figure 1C**).

**Figure 1:**
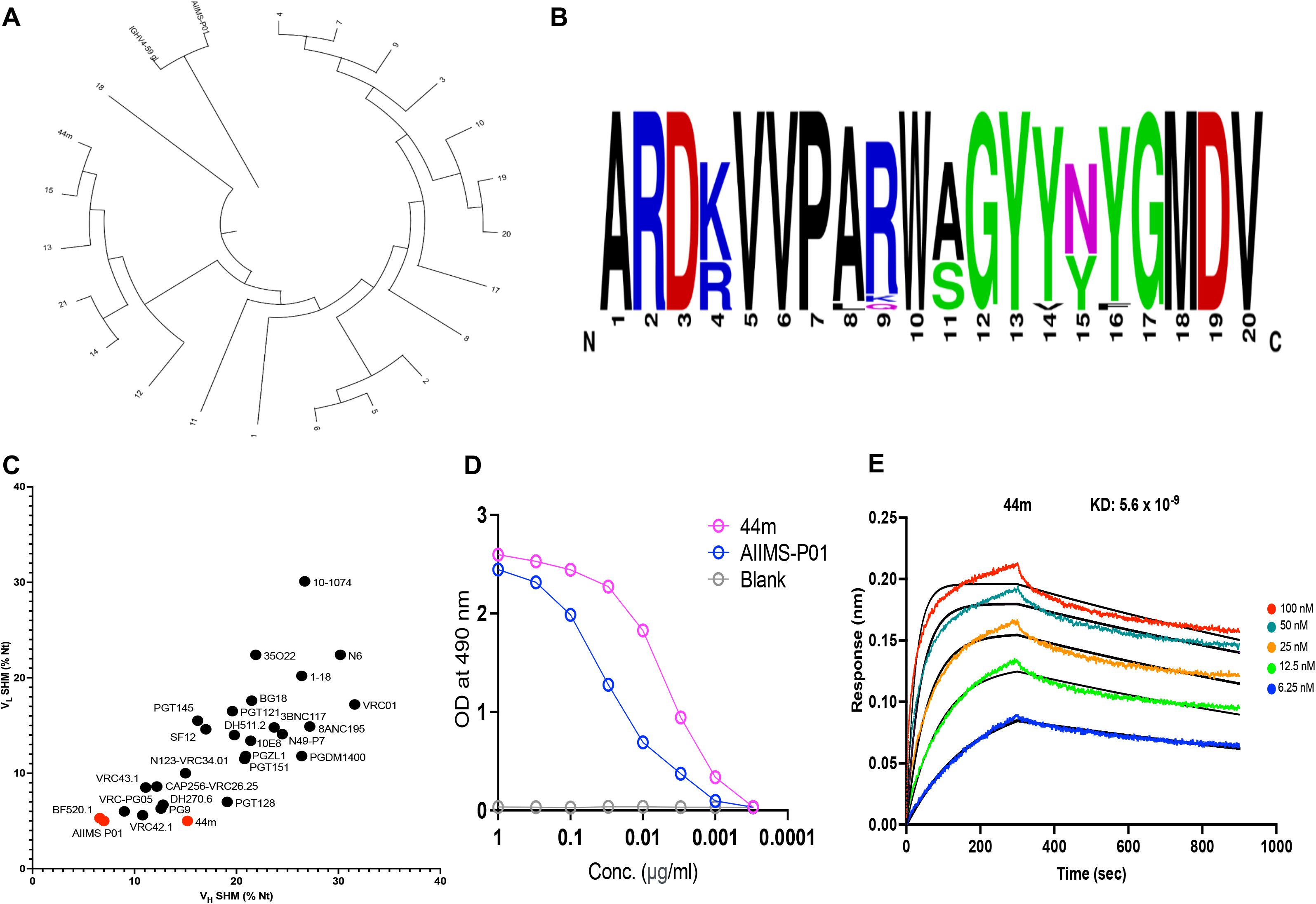
Characterization of AIIMS-P01 lineage members/clonotypes. **(A)** Phylogenetic tree of AIIMS-P01 lineage based on heavy chain sequences. **(B)** Logogram representing the frequency of the amino acid residues in CDRH3 of lineage mAbs. **(C)** Somatic hypermutation (SHM) analysis of HIV-1 adult and pediatric bnAbs shown as % nucleotide (Nt) mutations relative to respective germline variable gene sequence. Here, pediatric HIV-1 bnAbs are highlighted in red color. **(D)** Binding reactivity of the 44m and AIIMS-P01 mAbs to HIV-1 BG505.SOSIP.664.T332N gp140 trimeric envelope protein determined by ELISA. **(E)** Binding Affinity of 44m to BG505.SOSIP.664.T332N gp140 envelope trimer determined by Octet BLI. See also **Figure S1, S2 and Table S1 and S2**.

### Matured antibody 44m showed broader and more potent HIV-1 neutralization than AIIMS-P01

Next, we assessed the binding reactivity of the matured 44m mAb to heterologous HIV-1 BG505.SOSIP.664.T332N gp140 trimeric envelope protein in comparison to AIIMS-P01 WT and observed high binding efficiency (**Figure 1D**). To further validate the binding data obtained by ELISA, affinity analysis of the 44m with HIV-1 BG505.SOSIP.664.T332N gp140 trimer was performed using Octet BLI assays. Antibody 44m showed high nanomolar (nM) affinity (KD: 0.56 nM) with the BG505.SOSIP.664.T332N gp140 envelope trimer (**Figure 1E**). The neutralization potential of 44m antibody was tested against heterologous viruses and global panel of HIV-1 viruses, at concentrations ranging from 10µg/ml to 0.001 µg/ml, using a TZM-bl based neutralization assay^38^. The 44m antibody neutralized 77% HIV-1 clade-C viruses and 80% clade-B viruses and demonstrated an overall improvement of breadth of 79% against the heterologous viruses tested, and potency, with a geometric mean IC_50_ titer of 0.36 µg/ml, as compared to 67% exhibited by AIIMS-P01 previously^22^. Further, a 58% breadth, with IC_50_ titer of 0.43 µg/ml was observed, on testing this bnAb against the global panel of viruses (**Figure 2A and 2B**). The neutralizing activity data reveal an increase in potency and breadth of the matured version 44m against the heterologous viruses tested (**Figure 2C**). These findings encouraged us to perform structural characterization of 44m antibody with the stabilized envelope BG505.SOSIP.664.T332N gp140 trimer.

**Figure 2:**
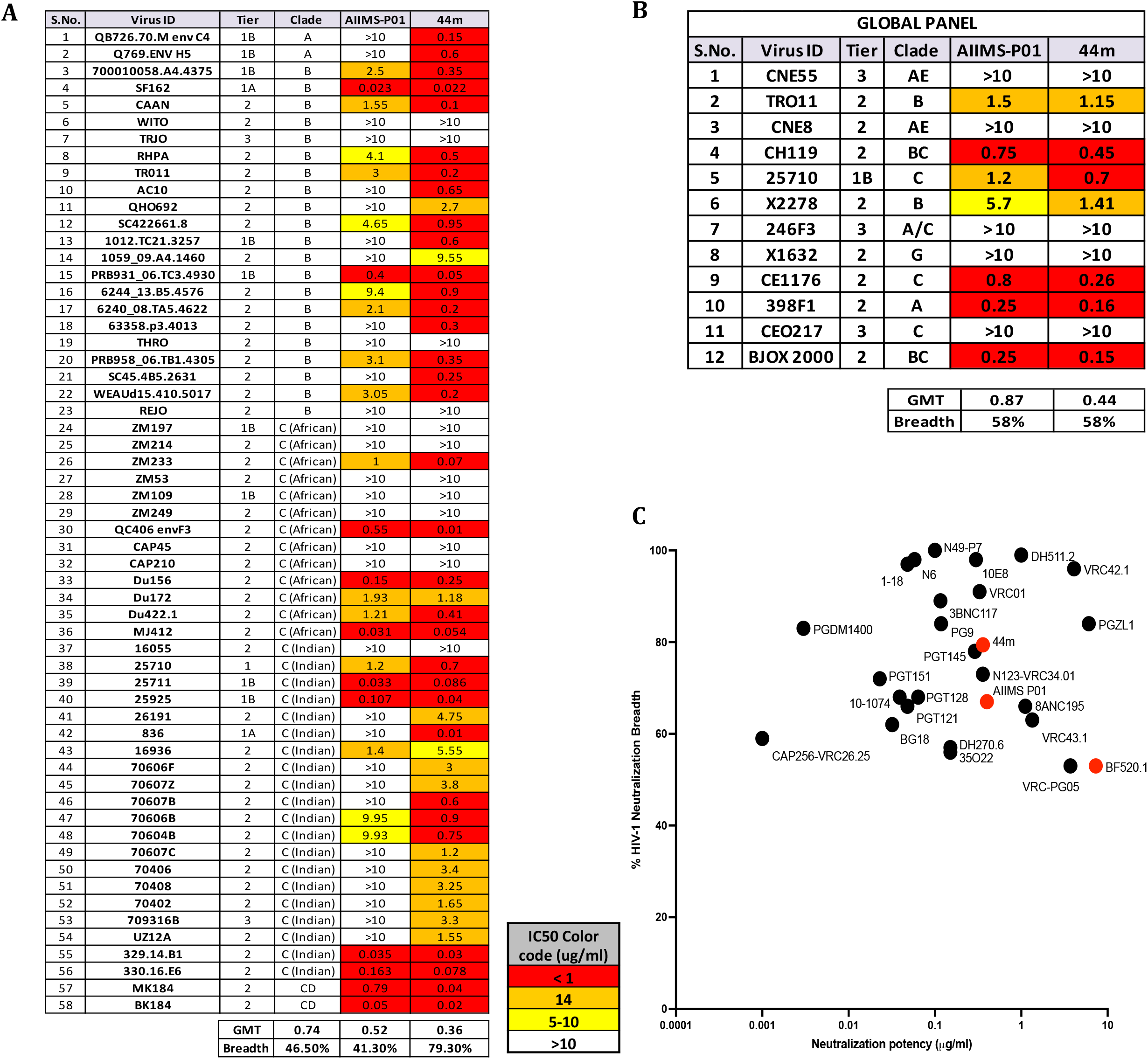
Matured AIIMS-P01 lineage antibody 44m showed broad HIV-1 neutralization with improved potency. **(A)** Heat map depicting IC_50_ values of the 44m tested against heterologous panel of HIV-1 viruses, using a TZM-bl based neutralization assay (n=58). **(B)** Heat map depicting IC_50_ values of the matured 44m mAb tested against HIV-1 global panel. **(C)** Neutralization breadth comparison of HIV-1 adult and pediatric bnAbs is shown. Here, pediatric HIV-1 bnAbs are highlighted in red color. The graph was plotted using Prism software. The neutralization potency (IC_50_) of 44m bnAb is compared with the IC_50_ of other HIV-1 bnAbs as documented in the Los Alamos HIV-1 molecular immunology database.

### Cryo-EM based structural analysis of 44m in complex with BG505.SOSIP.664.T332N gp140 trimer

A structural insight into mature 44m antibody in complex with BG505.SOSIP.664.T332N gp40 trimer was achieved through single particle cryo-Electron Microscopy (cryo-EM) and a high-resolution structure was solved at 4.4 Å resolution (**Figure 3 and S3–S6**). The atomic model fitted in the EM map of BG505.SOSIP.664.T332N gp140 trimer in complex with Fab of the 44m bnAb displayed a total of three 44m Fabs, with one Fab bound to each protomer of the Env trimer for effective neutralization (**Figure 3 and S3**). Cryo-EM structures of BG505.SOSIP.664.T332N gp140 Env trimer with 44m bnAb indicated trimeric shapes of HIV trimer, which additionally connected with extra densities, attributed to the corresponding bound Fab moieties (**Figure S4B**). Different subdomains, gp120, gp41 and Fab densities were visible in the high-resolution cryo-EM structure of the Env trimer in complex with 44m bnAb (**Figure 3**).

**Figure 3:**
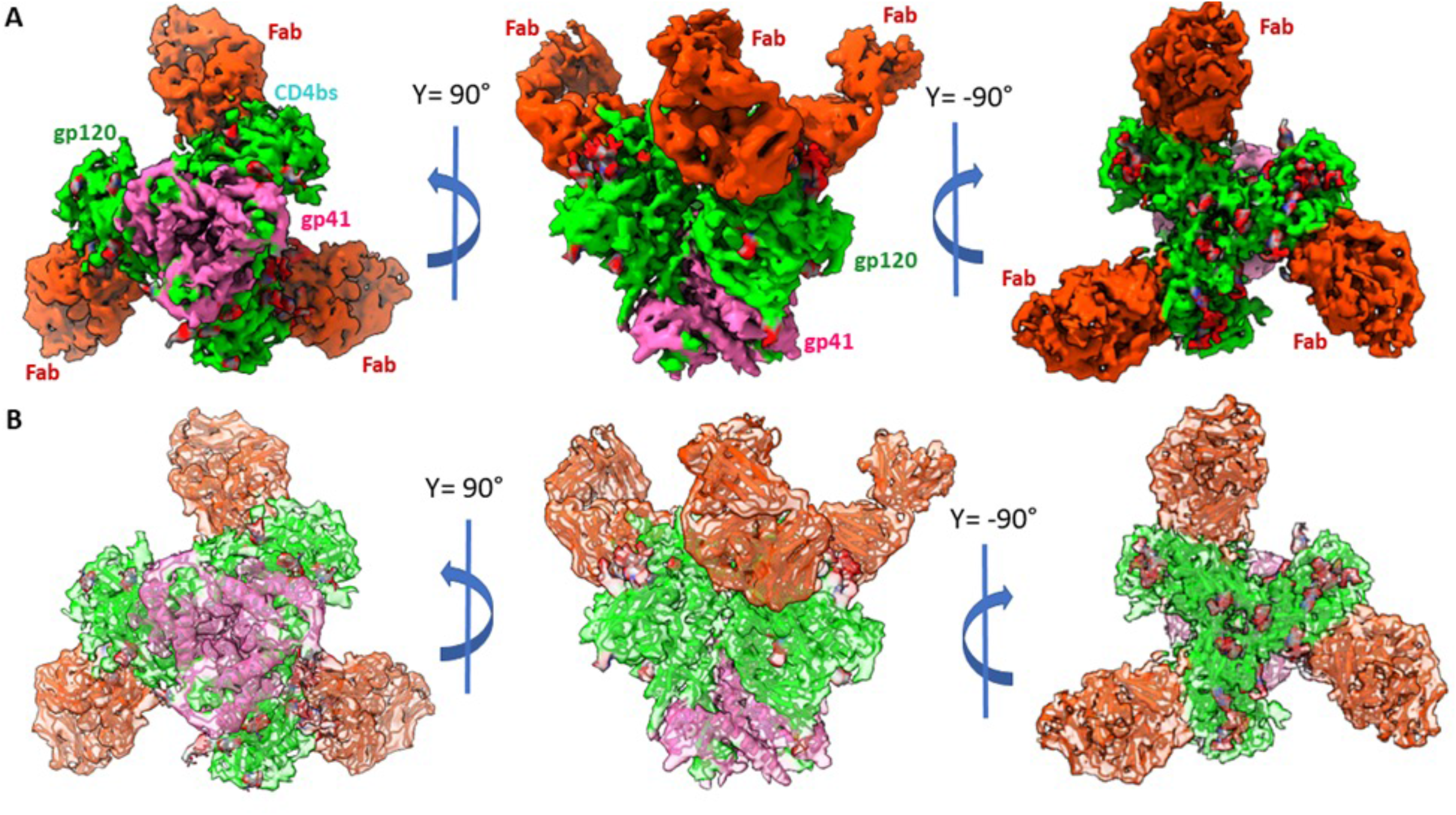
Cryo-EM reconstruction and model of BG505.SOSIP.664.T332N gp140 trimer in complex with 44m bnAb. **(A)** Side and top views of cryo-EM reconstructed map of BG505.SOSIP.664.T332N gp140 trimer in complex with 44m bnAb solved at ∼4.4Å resolution, with a local resolution in between 3.36 and 3.74Å. Color coding corresponding to segmented EM densities are: orange red, 44m bnAb; lime green, gp120; pink, gp41. **(B)** The corresponding atomic model fitted in the EM map of BG505.SOSIP.664.T332N gp140 trimer in complex with 44m bnAb showing three 44m bnAb binds to each gp120 monomeric subunit for effective neutralization. See also **Figure S3 – S6 and Table S3**.

Structural analysis revealed that the CDRH3 loop of neutralizing antibody 44m stretched deep inside the groove between the N295 and N332 residues, thus reaching the base of the V3 loop of gp120 and established a range of chemical bonds with the nearby favorable amino acids (**Figure 4A**). The total occupied surface area of 44m is ∼750 Å^2^, with extensive participation of the heavy chain. This surface area includes all the CDR regions and FR3 region, but the CDRH1 and CDRH3 loop show a predomination contribution. The positioning of the CDR regions of the heavy chain is crucial to establish contact points with the V3 loop of the gp120 (**Figure 4A, B**). The key residues on the CDRH3 loop of 44m mAb that are interacting with the gp120 region are V106, P107, A108, R109, and W110, whereas the primary contact points of CDRH1 and CDRH2 loops with the V3-region are H31, Y52, Y53, T54, D56, and T57 (**Figure 4B**). These interactions are mainly stabilized by both hydrogen bonding and van der Waals interactions. Interestingly, the R109 residue of the CDRH3 demonstrated a salt bridge interaction with the D325 residue of GDIR motif of the envelope gp120 region (**Figure 4C**). These interacting residues are hidden within the different clefts of the V3 loop of the gp120 region. The surface potential map of the paratope region depicts the interacting residues of CDR regions placed near the polar residues of the V3 loop (**Figure 4D and S7**).

**Figure 4:**
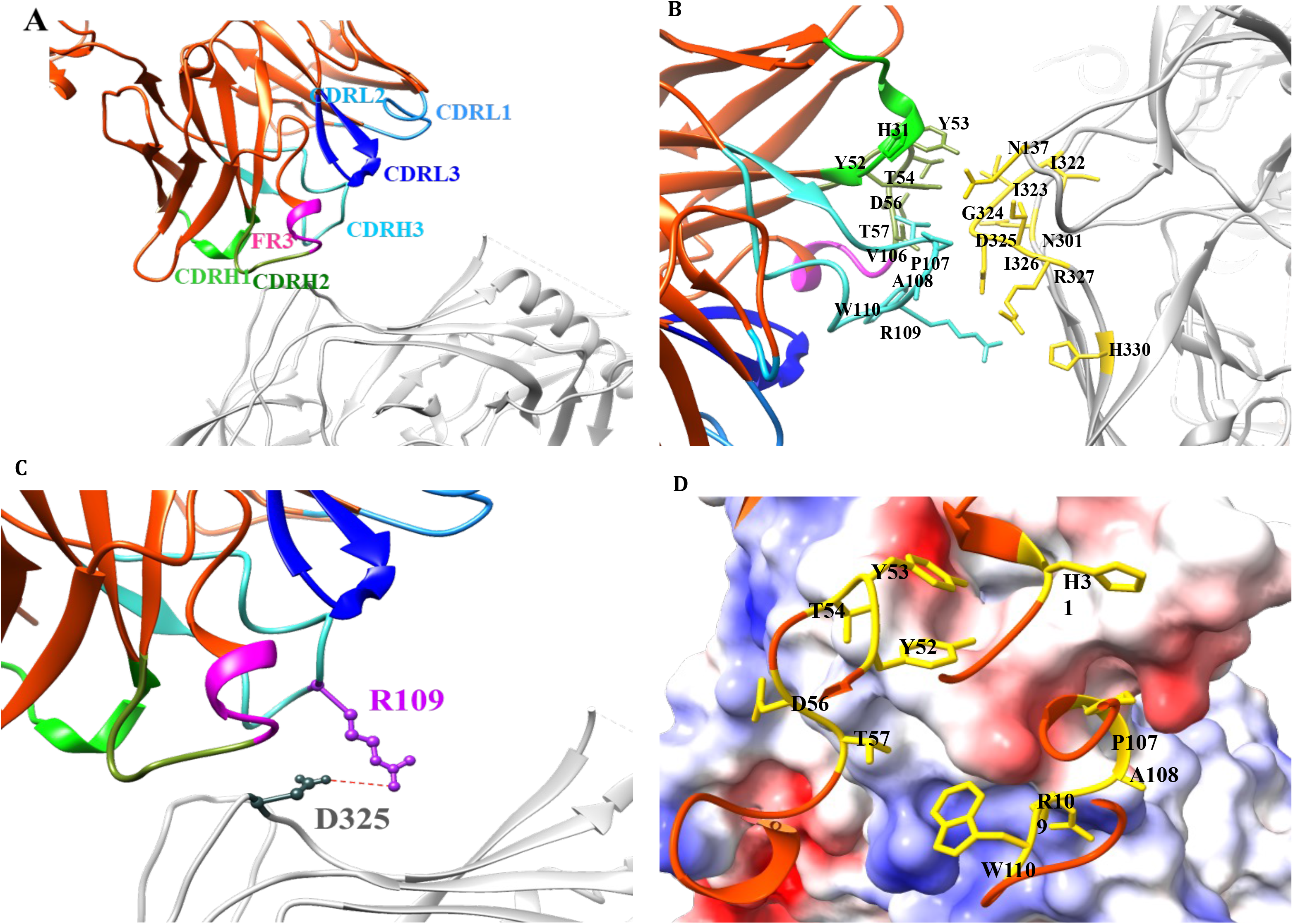
Interaction sites for 44m bnAb to form a complex with BG505.SOSIP.664.T332N gp140 trimer. **(A)** The CDR regions in the interface of gp120 and 44m bnAb are highlighted using color coding: magenta, FR3; green, CDRH1, olive green, CDRH2; and, cyan, CDRH3; dodger blue, CDRL1; deep sky blue, CDRL2; median blue, CDRL3. **(B)** Salt bridge interaction in between D325 residue in the V3 loop of envelope gp120 monomer and CDRH1 region, where D325 is shown in dark green and R109 is shown in purple color, and the bond is shown in red color. **(C)** Different interacting partners in the epitope region to adapt a stable conformational state with the CDR regions. **(D)** Electrostatic potential surface map of the interacting region of gp120 showing the interacting residues of the paratope facing towards the charged residues of the epitope. The residues on 44m bnAb is shown in golden color. See also **Figure S7 – S11**.

### Pediatric HIV-1 bnAb 44m primarily recognizes V3-glycans

The antibody-Env protein interaction studies unraveled the possibilities to explore the glycan antibody interactions. The atomic model demonstrated that the interaction sites for 44m bnAb on gp120 were positioned in an area that is surrounded by the N-glycan patches: Asn301, Asn332 located near the base of V3, Asn156 glycan in the V1V2 region, and Asn295 glycan near the bottom of gp120 (**Figure 5A-D**), though no interactions were observed with Asn156 and Asn295 glycans. This binding approach created an interlocked system of V_H_ loop of 44m and V3 loops of gp120, and forming a stable antibody-antigen complex (**Figure 5B**). Glycan binding to N301 is forming contacts with polar and charged amino acids of the CDRH2 loop, wherein the T54 and D56 of the CDRH2 region are interacting with the polar side of glycan moieties to mask the binding to host cells (**Figure 5C**). The N332 glycan engages in forming the largest interacting area, with most of the CDRH3 region involving different portions of glycan moieties on it. N-Acetyl glucosamine (NAG) attached to the N332 residue and forms a hydrogen bonding with Ser111 and Tyr113 residues of the CDRH3 region (**Figure 5D**). Overall, the structural analysis showed that the 44m antibody interacts with different glycan moieties attached to asparagine residues at 301 and 332 positions in the V3 region.

**Figure 5:**
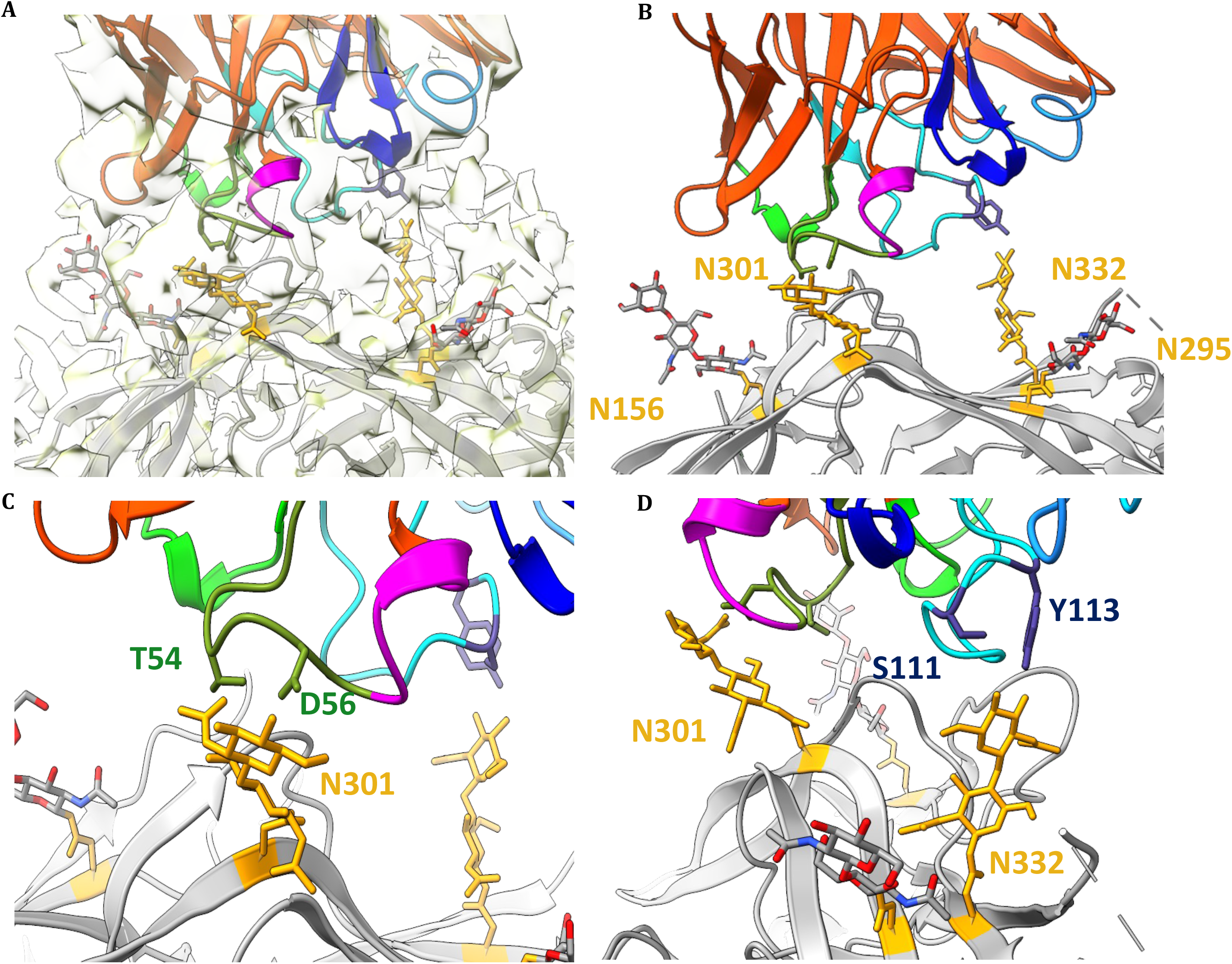
Glycan interaction sites of BG505.SOSIP.664.T332N gp140 trimer in complex with 44m Fab. **(A)** Positions and fitting different glycan residues are pointed in the EM map shown. In the atomic model N301 and N332 glycans are shown in orange color, whereas the Fab and CDR regions are colored similarly to Figure 2. **(B)** The gp120 monomer is shown in gray. Atomic model illustrating various glycan residues interacting with different region of CDR regions of 44m bnAb, where glycan attached to N156 is coming closer to CDRH1 region. **(C)** The NAG attached to Asn301 is making contact points with Thr54 and Asp56 in the CDRH2 region (shown in olive drab color). **(D)** Glycan binding to Asn332 is interacting with Ser111 and Tyr113 residues of the CDRH3 region (shown in deep blue color). See also **Figure S8 – S11**.

### 44m showed distinct glycan binding interactions, with similar pattern of epitope interaction like other V3-glycan bnAbs

A comparative structural analysis of the 44m antibody with other available bnAbs (BG18, 10-1074, DH270.6 & BF520.1) was performed to understand the exceptionality of this 44m antibody (**Figure S7 – S9**). Superimposition of the antibodies directed at the V3 loop region of the gp120 protomer showed a common interaction pattern across all the bnAbs within the V3 region. In the BG18 bnAb (6dfg) CDRL3 and CDRH3 loops are interacting with V3 region for stalking of the antibody onto the gp120 (**Figure S9A**). In bnAb 10-1074 (6udj), the V3 loop of gp120 protomer interacts with CDRH3 and CDRL3 regions (**Figure S9B**). DH270.6 (6um6) and BF520.1 (6mn7) have more inter-facial area, which helps in the interaction of residues in CDRL1 and CDRL2 regions. Additionally, the V1 region interacts with the CDRL3 loop (**Figure S9C, D**). The V3 loop has stable bonding with the CDRH1 loop for effective neutralization. The presence of an 8 amino acid long elongated face in the CDRH3 region in PGT121 and PGT122 classes of antibodies make these antibodies potent to bind gp120 surface using two functional surfaces (**Figure S9E**). These results suggest that the overall binding region of the CDR with both V3 and V1/V2 loops were significantly increased. The superimposition of V3-glycan bnAbs into the gp120 trimer depicts the variability in the binding surface area but a common binding pattern on the gp120.

### 44m HIV-1 bnAb showed co-dependence on GDIR motif and N332-supersite

Results obtained from structural mapping of 44m showed that this bnAb binds the N301, D325 residue of GDIR motif and N332-supersite env regions (**Figure 4 and 5**). Next, we used this structural mapping information to understand the linkage of these residues in 44m mediated HIV-1 neutralization by performing sequence alignment of these identified contact residues within the envelope regions of the viruses tested (**Figure S10**). The analysis revealed that the N156 and N301 glycans are relatively conserved among the 44m susceptible viruses (**Figure S10**). In contrast, mutations present at D325 and N332 positions were found to be associated with abrogation in neutralization potential of 44m and vice-versa, e.g. HXB2, ZM249 and ZM233 viruses were resistant to neutralization by 44m, due to the absence of GDIR epitope in these viruses. To further confirm the reliance of the 44m mAb on the D325 residue, we conducted functional mutant assays and noted a 59-fold decrease in the presence of the D325K mutation, highlighting the dependence on the GDIR motif (**Figure S11**). These findings suggest that the neutralization dependence of the pediatric bnAb 44m relies on the D325 residue of GDIR motif and N332-spersite, as has also been reported for the adult HIV-1 V3-glycan directed bnAbs PGT121 and 10-1074^26, 39^.

## DISCUSSION

The footprints of HIV-1 broadly neutralizing antibodies (bnAbs) can provide a template for structure-guided vaccine design^6, 7^. Highly potent bnAb based therapeutics, prophylactics and vaccines are attractive strategies to tackle HIV-1^3^. Immunogenetics based information of potent HIV-1 bnAbs derived from deep sequencing or single cell analysis of B cell repertoire (BCR) of infected donors can provide critical insights towards understanding their natural development during the course of infection and reveal the frequency of B cells within the human B cell repertoire, that can elicit potent bnAbs to guide vaccination strategies^6, 29, 40, 41^. Further, the structural characterization of potent HIV-1 bnAbs in complex with the viral envelope provides useful mechanistic insights of viral neutralization and information of epitope-paratope interaction for rational vaccine design^7^. Currently, the leading strategy in innovative HIV-1 next-generation immunotherapeutic and vaccine design is to develop and elicit bnAb responses by steering bnAb expressing B cells^6, 8^. To achieve this goal, it is essential to understand the structural mechanism of HIV-1 neutralization and immunogenetics of B cells that elicit potent bnAbs. The evolution of HIV-1 bnAbs from adult donors has been studied extensively^29, 42, 43^, but, no information is available on the evolving HIV-1 bnAb lineage in chronically infected children.

Herein, to fill this knowledge gap, we synthesized a heavy chain matured lineage member (44m) of AIIMS-P01 bnAb identified from our recent study on identification of multiple epitopes targeting adult HIV-1 bnAbs clonotypes in AIIMS_330 pediatric elite-neutralizer from longitudinal bulk B cell repertoire analysis by NGS^35^. We further evaluated the structural features and functionality of 44m antibody in terms of viral binding and neutralization activity. We used only heavy chain of AIIMS-P01 matured lineages as we didn’t find matched CDRL3 of the AIIMS-P01 light chain in our deep sequencing data, plausibly due to the low depth of sequencing^35^. We combined functional and structural approaches and showed that maturation in the 44m heavy chain, like that reported for the adult bnAbs of the PGT class^42, 44^, was functionally important for HIV-1 Env binding and neutralization. This was demonstrated by the increased heterologous breadth observed when the mature heavy chain was paired with the original light chain of AIIMS-P01 WT bnAb, suggesting that the recently acquired SHMs can be functionally important for the evolution and neutralization breadth for this bnAb. In addition, the matured bnAb 44m exhibited a change of indel sequence (SNPSR mutated to SDPIR), in comparison to its WT bnAb AIIMS-P01 (**Figure S1**).

Unlike HIV-1 CD4bs bnAbs, the V3-glycan targeting bnAbs are of high interest because they are common and not restricted by certain germline genes^1, 3^. The pediatric bnAb AIIMS-P01 and infant derived BF520.1 are of particular interest because these showed broad HIV-1 neutralization despite limited SHM^10, 22^. The 44m lineage bnAb identified and characterized herein showed improved neutralization potency and breadth plausibly by acquisition of higher number of SHM, that led to an increase in potency twice that of the WT AIIMS-P01 bnAb. To estimate the probability of amino acid SHM substitutions in the 44m heavy chain we used the ARMADiLLO method as described previously . Interestingly, we identified 4 improbable AA SHM in 44m bnAb with less than 2% frequency. The probability of antibody mutations can be useful to vaccinologists in designing vaccines to elicit such bnAbs which are enriched with developmentally rate-limiting improbable mutations^36, 37^.

The cryo-EM structural analysis of bnAb 44m, indicated that CDRH1 (H31, Y53 and T57) and CDRH3 (P107, A108 and R109) residues appear to contribute to the 44m paratope by mediating contacts with the conserved V3-glycan N332-supersite, D325 of ‘GDIR’ sequence motif, glycans at position 301 and 332. These major determinants of neutralization breadth interacting with residues within the CDRH1 and CDRH3 regions of the 44m, are similar to that in the adult bnAbs and distinct from the infant bnAb BF520.1^10, 42^. The angle of approach by 44m towards the V3 epitope was previously determined to be similar to PGT121^39, 45^, although the positioning of 44m was notably different and slightly rotated relative to the PGT121. The crystal structure of PGT121 in complex with gp120 identified the GDIR motif and glycans at positions N332 and N301 as the primary contacts defining the PGT121 epitope^39, 45^. The CDRH3 loop, which is highly mutated in PGT121, penetrates the glycan shield in order to contact both the GDIR motif and N332 glycan.

The structural model similarly indicates potential CDRH3 contacts with the GDIR motif and N332 glycan. However, structurally defined epitope-paratope interface do not fully capture the functional binding contacts that drive neutralization and escape^46^, reinforcing the gravity of functional assays to define the recognition determinants that are important for neutralization activity. As we demonstrated previously the WT mAb AIIMS-P01 does not show the dependence on V1 glycan N156^22^, likewise the structural analysis of mutant 44m suggests N156 is hanging out of the core, without making dominant interactions. Though we observed that 44m epitope is surrounded with other V3-region glycans including N295 and N301, however, HIV-1 viral sequence alignment (**Figure S10**) revealed that 44m is primarily dependent on N332-glycan supersite and GDIR motifs. To further validate the 44m mAb dependence on D325 residue, we performed functional mutant assays and observed a 59-fold decrease in case of D325K mutation indicating dependence on GDIR motif (**Figure S11**). Based on the findings of HIV-1 binding, neutralization and structural analysis of the 44m pediatric bnAb, we postulate that a germline targeting vaccine / immunogen could potentially elicit AIIMS-P01 / 44m-like responses. In the AIIMS_330 pediatric elite neutralizer, the circulating and coevolving viruses may have led to elicitation of this bnAb lineage (44m), to drive affinity maturation in the AIIMS-P01 bnAb, as has been observed previously for the evolution of V1V2 and V3-glycan plasma bnAbs^20^.

In summary, the structural and functional characterization of a heavy chain matured pediatric bnAb 44m showed improved HIV-1 neutralization potency and breadth in comparison to its WT bnAb AIIMS-P01. This study for the first time provides evidence towards contribution of antibody SHM in improved HIV-1 neutralizing efficiency of a bnAb identified from a pediatric donor living with chronic HIV-1 clade C infection. Further studies in this direction are required to be conducted to understand the antigenic triggers in chronically infected children that can elicit similar protective bnAbs targeting other HIV-1 bnAb epitopes which in turn can provide a blueprint to guide HIV-1 vaccine design.

### Limitations of the study

Our work is focused on the binding, neutralization, and structural characterization of a matured lineage member (44m) of AIIMS-P01 pediatric HIV-1 bnAb. All conclusions are based on cryo-EM structural analysis, genetic features analysis and in-vitro HIV-1 viral neutralization assays. The bulk B cell repertoire of total PBMCs led to identification of only few AIIMS-P01 heavy chain lineage members/clonotypes and no light chain sequences were identified. An in-depth NGS based sequencing of AIIMS-P01 lineage will enable the identification of a large number of clonotypes for in depth characterization of bnAb lineages.

## STAR ★ METHODS

Detailed methods are provided in the online version of this paper and include the following:

- **KEY RESOURCES TABLE**
- **RESOURCE AVAILABILITY**

`o` Lead contact
`o` Materials availability
`o` Data and code availability
- **EXPERIMENTAL MODEL AND STUDY PARTICIPANT DETAILS**

`o` Cell Lines
- **METHOD DETAILS**

`o` Identification of AIIMS-P01 lineage members
`o` 44m heavy chain SHM analysis using ARMADiLLO method
`o` Antibody gene synthesis
`o` Antibody genes sequence analysis
`o` Expression of monoclonal antibodies
`o` Expression and purification of HIV-1 trimeric proteins
`o` Binding analysis of mAbs by ELISA
`o` HIV-1 pseudovirus generation
`o` HIV-1 neutralization assays
`o` Fab Fragment Preparation
`o` Octet BLI analysis
`o` Negative-stain EM
`o` Sample preparation for cryoEM
`o` CryoEM data acquisition
`o` CryoEM data analysis and model building
- **QUANTIFICATION AND STATISTICAL ANALYSIS**

## KEY RESOURCES TABLE

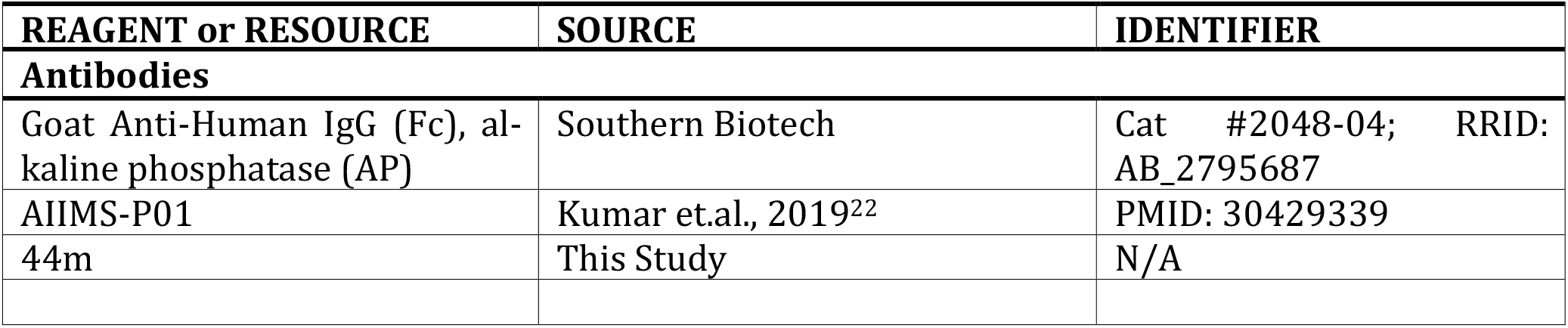
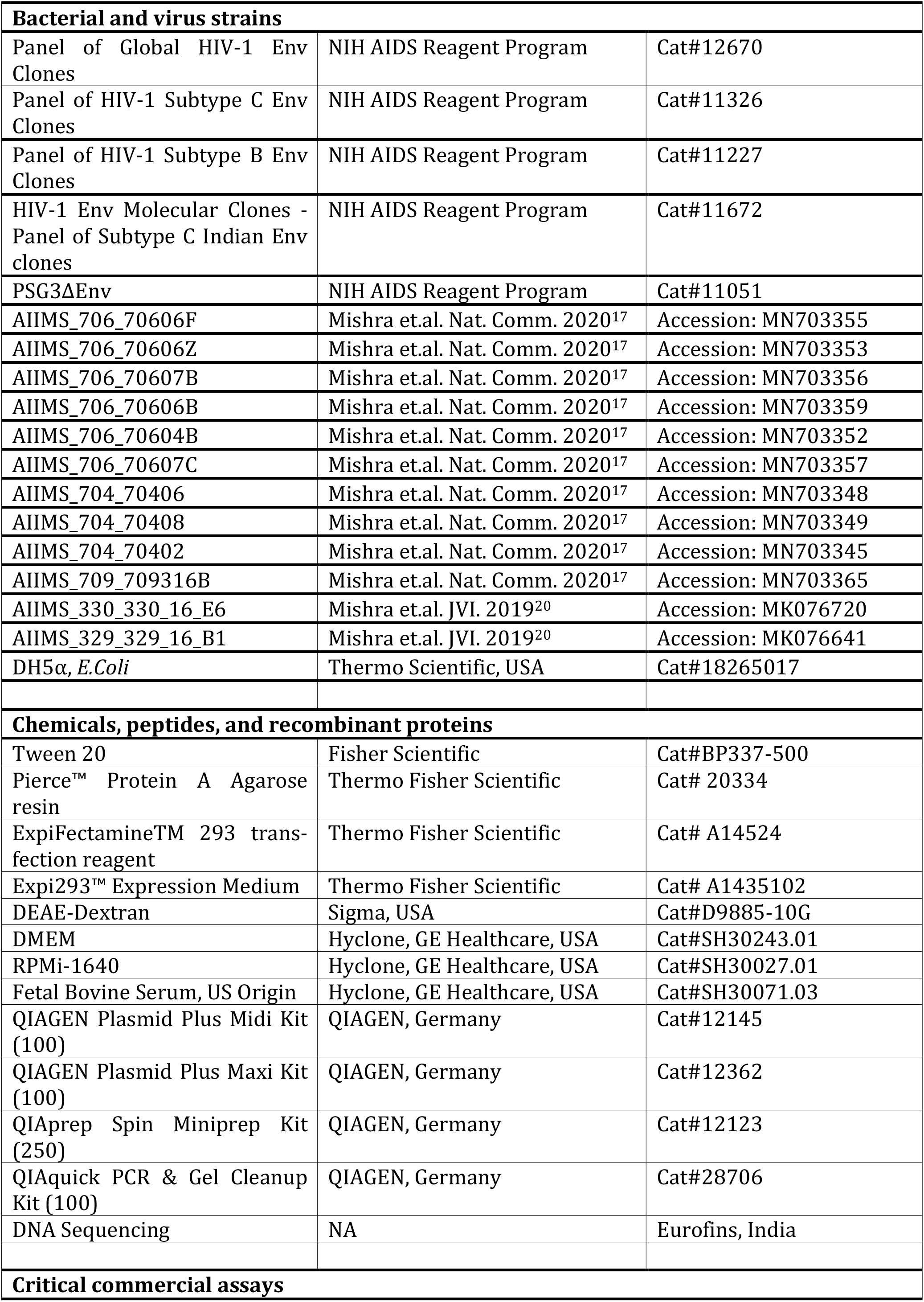
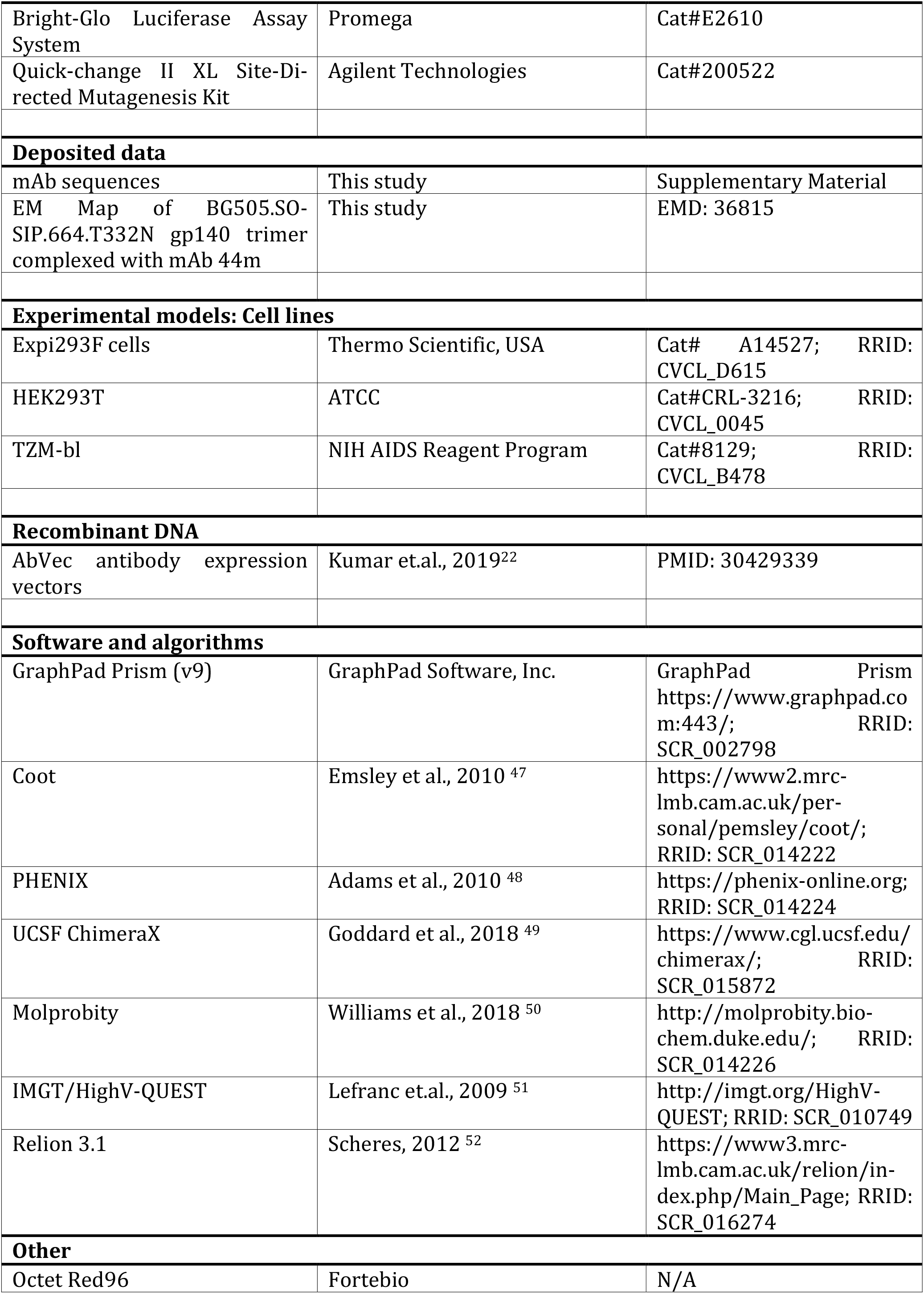

## RESOURCE AVAILABILITY

### Lead contact

Further inquiries and requests for data, plasmids and resources should be directed to the lead contact Kalpana Luthra.

### Materials availability

Antibody expression plasmids generated in this study (see key resources table) are available upon request from the lead contact with a completed Materials Transfer Agreement.

### Data and code availability

- Atomic coordinates and cryo-EM maps for reported structures are deposited into the Elec-tron Microscopy Data Bank (EMDB) with accession code EMD-36815 for BG505.SO-SIP.664.T332N gp140 envelope trimer in complex with mAb 44m. The sequences of the 44m and AIIMS-P01 lineage antibody heavy chain variable regions are present in the sup-plementary materials. Raw sequence data that support the findings in this study has been deposited at NCBI Sequencing Read Archive (www.ncbi.nlm.nih.gov/sra) and publicly available under BioProject accession number SRA: PRJNA999025. Processed datasets are available at https://github.com/prashantbajpai/HIV_BCR_Analysis.
- All code for analysis and generating individual figure panels are available on GitHub and link to the repository is https://github.com/prashantbajpai/HIV_BCR_Analysis.
- Any additional information required to reanalyze the data reported in this paper is availa-ble from the lead contact upon request.

## EXPERIMENTAL MODEL AND STUDY PARTICIPANT DETAILS

### Cell Lines

Human embryonic kidney (HEK)-derived 293T, and HeLa-derived TZM-bl cells were main-tained in complete Dulbecco’s Modified Eagle Medium (herein referred to as cDMEM) contain-ing high glucose Dulbecco’s Modified Eagle Medium (DMEM, Thermo Fisher), 1X Penicillin-Streptomycin (Pen Strep, Thermo Fisher) and 10% fetal bovine serum (FBS, Gibco) at 37⁰C and 5% CO_2_. FreeStyle 293F and Expi293F cells (Thermo Fisher) were maintained in Freestyle 293 Expression Medium and Expi293 Expression Medium, respectively, at 37⁰C and 8 % CO2 with shaking at 120 RPM.

## METHOD DETAILS

### Identification of AIIMS-P01 lineage members

We recently reported the deep sequencing of bulk B cell repertoire (BCR) that was performed using primers, protocols and Illumina MiSeq (MiSeq Reagent Kit v3, 600-cycle) as described previously^35, 53^. The Abstar analysis pipeline was used as previously described to quality trim, remove adapters and merge paired sequences^53^. Sequences were then annotated with Abstar in combination with UMI based error correction by AbCorrect (https://github.com/briney/abtools/). For comparison of frequencies, read counts were scaled for each repertoire as previously described due to the large differences in the number of reads between each group. Somatic hypermutation (SHM) was calculated using the R package Shazam^54^. Clonotype analysis was performed using Immcantation pipeline^54, 55^. Sequences were grouped into clonotypes based on nucleotide hamming distance of 0.16 calculated based on bimodal distribution of distance of each sequence with its nearest neighbor. Alternatively, sequences were also clustered into clonal groups using an in-house script. The criteria used for clonal assignment was sequences having same V and J gene usage, same CDRH3 length and at least 80% CDRH3 amino acid identity. Germline V(D)J sequence was reconstructed using IMGT-gapped reference V, D and J sequences.

### 44m heavy chain SHM analysis using ARMADiLLO method

To estimate the probability of amino acid SHM substitutions in the 44m heavy chain we used the ARMADiLLO method as described previously^36^.

### Antibody gene synthesis

The antibody heavy chain genes of matured AIIMS-P01 lineage antibody was synthesized after codon-optimization for mammalian expression from Genscript, Inc. USA, and cloned in respective monoclonal antibody expression vector AbVec under AgeI and SalI sites^56^.

### Antibody genes sequence analysis

The sequencing of the antibody genes was done commercially from Eurofins, India. The sequences were analyzed online through IMGT/V-QUEST (http://www.imgt.org/IMGT_vquest/vquest)^51^.

### Expression of monoclonal antibodies

All HIV-1 mAbs were expressed in Expi293F cells (Thermo Fisher) as described previously^22, 57^. Briefly, 15µg each of heavy chain and light chain expressing IgG1 plasmids were co-transfected using PEI-Max as transfection reagent. Following 4-6 days of incubation, cells were harvested by centrifugation and filtered through 0.22 mm syringe filter (mdi). The supernatant was added to a Protein A column affinity chromatography column (Pierce). The column was then washed with 1×PBS and mAbs were eluted with IgG Elution Buffer (Pierce), immediately neutralized with 1M Tris pH 8.0 buffer and extensively dialyzed against 1×PBS at 4°C. The mAbs were then concentrated using 10kDa Amicon Ultra-15 centrifugal filter units (EMD Millipore), filtered through a 0.22 mm syringe filter (mdi) and stored at -80°C for further use.

### Expression and purification of HIV-1 trimeric proteins

The BG505.SOSIP.664.T332N gp140 trimeric proteins with twin-strep-tag was expressed in HEK 293F cells and purified by methods described previously^58^. Briefly, SOSIP proteins were expressed in transiently transfected HEK293F suspension cells (Invitrogen, cat no. R79009), maintained in FreeStyle Expression Medium (Gibco). For transfection, HIV-1 Env and furin protease-encoding plasmids were mixed in a 3:1 Env to furin ratio (w/w) were incubated with PEImax (Polysciences Europe GmBH, Eppelheim, Germany) in a 3:1 (w/w) PEImax to DNA ratio and then added in the supernatant of cells at a density of 1.2-1.5 million cells/mL. Six days post-transfection, supernatants were harvested, centrifuged, and filtered using 0.22 µm pore size filters before protein purification. HIV-1 env proteins were purified by immunoaffinity chromatography with PGT145 antibody affinity column. Unpurified proteins contained in HEK293F filtered supernatants were captured on PGT145-functionalized CNBr-activated sepharose 4B beads (GE Healthcare) by overnight rolling incubation at 4 °C. Subsequently, the mixes of supernatant and beads were passed over Econo-Column chromatography columns (Biorad). The column was then washed with three column volumes of a 0.5 M NaCl and 20 mM Tris HCl pH 8.0 solution. After elution with 3 M MgCl_2_ pH 7.4, proteins were buffer-exchanged into TN75 (75 mM NaCl, 20 mM Tris HCl pH 8.0) buffer by ultrafiltration with Vivaspin20 filters (Sartorius) of MWCO 100 kDa. Protein concentrations were determined from the A280 values measured on a NanoDrop2000 device (Thermo Fisher Scientific) and the molecular weight and extinction coefficient values calculated by the ProtParam Expasy webtool. Purity was assessed by blue native polyacrylamide gel electrophoresis (BN-PAGE) and binding reactivity with HIV-1 bnAbs was assessed by ELISA.

### Binding analysis of mAbs by ELISA

Briefly, 96-well ELISA plates (Costar) were coated with 5µg/ml recombinant HIV-1 gp120 monomeric proteins overnight at 4°C in 0.1 M NaHCO_3_ (pH 9.6). Next day, plates were washed thrice with 1×PBS (phosphate buffered saline) and blocked with 15% FBS RPMI and 2% BSA. After 1.5 hours of blocking at 37°C, plates were washed thrice with 1×PBS. Then, serial dilutions of monoclonal antibodies (mAbs) were added and incubated for 1 hour at 37°C. Next, alkaline phosphatase (AP) labelled anti-Fc secondary antibody (Southern Biotech) at 1:2,000 was added and plates were incubated at 37°C for 1 hour. Plates were then washed thrice with 1×PBS and AP substrate tablets (Sigma) dissolved in diethanolamine (DAE) was added and incubated for 30 min at room temperature in the dark and readout was taken at 405nm. The BG505.SOSIP.664.T332N gp140 trimeric ELISA was performed as described previously^58^. Briefly, purified twin-strep-tag BG505.SOSIP.664.T332N gp140 protein (1 µg/mL) was diluted in PBS and captured on 96-well Streptactin XT ELISA plates (IBA, Germany) followed by a 2 h incubation at room temperature. Following two washes with 1x PBS to remove unbound trimers, serial dilutions of test antibodies in PBS/2% skimmed milk were added and incubated for 2 h. After 3 washes with PBS, alkaline phosphatase (AP) labelled anti-Fc secondary antibody (Southern Biotech) at 1:2,000 in PBS/2% skimmed milk was added and incubated for 1 h, followed by 4 washes with PBS/0.05% Tween20. Plates were then washed thrice with 1×PBS and AP substrate tablets (Sigma) dissolved in diethanolamine (DAE) was added and incubated for 30 min at room temperature in the dark and readout was taken at 405nm.

### HIV-1 pseudovirus generation

The HIV-1 pseudoviruses were produced in HEK 293T cells as described earlier^20, 22^ by co-transfecting the full HIV-1 gp160 envelope plasmid and a pSG3ΔEnv backbone plasmid. Briefly, 1×10^5^ cells in 2ml complete DMEM (10% fetal bovine serum (FBS) and 1% penicillin and streptomycin antibiotics) were seeded per well of a 6 well cell culture plate (Costar) the day prior to co-transfection for HIV-1 pseudovirus generation. For transfection, envelope (1.25µg) to delta envelope plasmid (2.50µg) ratio was 1:2, this complex was made in Opti-MEM (Gibco) with a final volume of 200µl for each well of the 6 well plate and incubated for 5 minutes at room temperature. Next, 3µl of PEI-Max transfection reagent (Polysciences) (1mg/ml) was added to this mixture, mixed well and further incubated for 15 min at room temperature. This mixture was then added dropwise to HEK 293T cells supplemented with fresh complete DMEM growth media and incubated at 37°C for 48 hours. Pseudoviruses were then harvested by filtering cell supernatants with 0.45 mm sterile filter (mdi) and stored frozen at −80°C as aliquots.

### HIV-1 neutralization assays

The HIV-1 neutralization assays of monoclonal antibodies (mAbs) were done as described earlier^38, 59^. Neutralization was measured as a reduction in luciferase gene expression after a single round of infection of TZM-bl cells (NIH AIDS Reagent Program) with HIV-1 envelope pseudoviruses. The TCID_50_ of the HIV-1 pseudoviruses was calculated and 200 TCID_50_ of the virus was used in neutralization assays by incubating with 1:3 serially diluted mAbs starting at 10 μg/ml. After that, freshly trypsinized TZM-bl cells in growth medium (complete DMEM with 10% FBS and 1% penicillin and streptomycin antibiotics) containing 50μg/ml DEAE Dextran and 1 mM Indinavir (in case of primary isolates) at 10^5^ cells/well were added and plates were incubated at 37°C for 48 hours. Virus controls (cells with HIV-1 virus only) and cell controls (cells without virus and antibody) were included. MuLV was used as a negative control. After the incubation of the plates for 48 hours, luciferase activity was measured using the Bright-Glow Luciferase Assay System (Promega). IC_50_ for antibodies were calculated. Values were derived from a dose-response curve fit with a non-linear function using the GraphPad Prism 9 software (San Diego, CA).

### Fab Fragment Preparation

The Fab fragments were generated from 4 mg of 44m IgG antibody using a Fab Fragmentation Kit (G Biosciences) according to manufacturer’s protocol. Purity and size of Fab fragments were assessed by SDS-PAGE.

### Octet BLI analysis

Octet biolayer interferometry (BLI) was performed using an Octet Red96 instrument (ForteBio, Inc.). A 5 μg/ml concentration of each mAb was captured on a protein A sensor and its binding kinetics were tested with serial 2-fold diluted HIV-1 SOSIP trimer protein (100 nM to 6.25 nM). The baseline was obtained by measurements taken for 60 s in BLI buffer (1x PBS and 0.05% Tween-20), and then, the sensors were subjected to association phase immersion for 300 s in wells containing serial dilutions of HIV-1 SOSIP protein. Then, the sensors were immersed in BLI buffer for as long as 600 s to measure the dissociation phase. The mean Kon, Koff and apparent KD values of the mAbs binding affinities for HIV-1 SOSIP envelope protein were calculated from all the binding curves based on their global fit to a 1:1 Langmuir binding model using Octet software version 12.0.

### Negative-stain EM

To observe the binding pattern of 44m bnAb to BG505.SOSIP.664.T332N gp140 trimer and the homogeneity of the complex, we first performed room temperature negative staining TEM. The SEC purified complex of BG505.SOSIP.664.T332N gp140 trimer and 44m bnAb (1.2 mg/ml) was diluted by 70 times for analysis. The 3.5 µl of sample mixture was put onto a glow discharged carbon coated Cu grids for 30 secs (EM grid, 300 mesh, Electron Microscopy Sciences). After 1.5 min of incubation of the sample on the grid, the remained solvent was blotted and three drops of 1% uranyl acetate (Uranyl Acetate 98%, ACS Reagent, Polysciences, Inc.) was applied on the grid for staining purpose. The excess stain was blotted after each addition and after air dried, the grid was used for data collection with 120 kV Talos L120C electron microscope. Data acquisition was performed using 4k × 4k Ceta camera at the magnification of 73kx and it is calibrated at 3.84Å/pixel. The collected images were processed in EMAN 2.1^60^. From these micrographs we picked particles in both manual and automated mode, and its co-ordinates were extracted using e2boxer.py in EMAN 2.1. Followed by, reference free 2D class averages were performed to analyze different views of bnAb bound trimer complex. The cleaned particles after extraction were taken for reference-free 2D class averages using simple_prime2D of SIMPLE 2.1 software^61^ with a mask diameter of 30 pixels at 3.84 Å/pix.

### Sample preparation for cryo-EM

R1.2/1.3 300 mesh copper grids (Quantifoil) (Electron Microscopy Sciences) were glow discharged at 20mA for 90 seconds before cryo-freezing. Three microliters of the SEC purified complex of BG505.SOSIP.664.T332N gp140 trimer and 44m bnAb (1.2 mg/ml) was applied onto the freshly glow discharged grid, and immediately blotted for 8.5 secs without any blot force just after 10 secs of incubation to remove excess solvent in pre-equilibrated chamber of FEI Vitrobot Mark IV plunger. The sample was plunged into the liquid ethane just after blotting.

### Cryo-EM data acquisition

Cryo-EM data were collected using 200 kV TalosArctica transmission electron microscope (Thermo Scientific™) equipped with Gatan K2 Summit Direct Electron Detector. Movies were recorded automatically using Latitude-S (DigitalMicrograph - GMS 3.5) at nominal magnification of 45,000x at the effective pixel size of 1.17 Å (*14*). Micrographs were acquired in counting mode with a total dose of 60 e^-^/Å2, with an exposure time of 8 sec distributed for 20 frames. A total of 3000 movies were acquired for the BG505.SOSIP.664.T332N gp140 trimer and 44m bnAbs protein complexes respectively.

### CryoEM data analysis and model building

Single-Particle Analysis (SPA) were performed for the acquired cryo-EM movies using the Relion version 3.1^52^. At first, drift and gain corrections of the individual movies were performed with MotionCorr2^62^ and estimated Contrast transfer function (CTF) parameters using CTFFIND 4.1.13^63^. Subsequently, CTF estimated micrographs were subjected to analyze to eliminate bad micrographs using cisTEM^64^ and also, to remove poor resolution micrographs with a fit resolution threshold of 7 Å. The particles from best micrographs were chosen for automated picking using 2D reference in Relion and extracted with the box sizes of 280 Å for the BG505.SOSIP.664.T332N gp140 trimer and 44m bnAb complexes. After three rounds of rigorous 2D classification good classes with high-resolution features of the complex were obtained as 1080743 particles. These well-defined particles were selected for 3D classification with C3 symmetry. To achieve high resolution, all particles belonging to the best classes of the complex was accounted for 3D auto-refinement and followed by movie refinement. The sharpening for the 3D auto-refined maps was performed with Relion 3.1^52^ and PHENIX^48^. Overviews of cryo-EM data processing is shown in **Table S3**. Global resolution of Fourier shell correlation (FSC) was estimated at the threshold of 0.143 and the estimation of local resolution were performed with ResMap, using two auto-refined half maps.

Automated model building was iteratively done with Phenix Real Space Refinement. Only the Env trimer (PDB ID: 5aco) was docked with cryo-EM maps using UCSF Chimera “Fit in map” tool. To build the model for bnAb, the query sequences of the Fab was submitted to Swiss-Model and the resultant model was also fitted in the EM maps. The structural statistics for cryo-EM map and atomic model were analyzed using Phenix^48^, EMringer^65^, Molprobity^50^, and UCSF chimera^66^. cryo-EM map and atomic model were visualized using UCSF ChimeraX^49^.

## QUANTIFICATION AND STATISTICAL ANALYSIS

For all HIV-1 pseudoviruses neutralization assays, data were fitted asymmetric nonlinear regression model to obtain the IC_50_. All neutralization assays were repeated at least 2 times, and data shown are from representative experiments. All statistical analysis was done with GraphPad Prism software version 9.

## ACKNOWLEDGMENTS

This antibody work was supported by Department of Biotechnology, India Indo-SA BT/PR2450/MED/29/1222/2017) grant awarded to K.L. The cryoEM work and consumables were supported by DBT BUILDER PROGRAM (BT/INF/22/SP22/844/2107), DST-FIST (SR/FST/LSII-039/2015) and SERB (SERB-EMR/2016/000608) grants awarded to S.D. S.K. is supported through DBT/Wellcome Trust India Alliance Early Career Fellowship grant IA/E/18/1/504307 (S.K.). We are very much thankful to NIH AIDS reagent program for HIV-1 research reagents, Neutralizing antibody consortium (NAC), IAVI, USA for HIV-1 neutralizing antibodies.

## AUTHOR CONTRIBUTIONS

Experimental work, data acquisition and analysis of data by S.Ku., S.D.S., A.C., P.B., S.S, S.Ka, R.L., S.D. Conceptualization and implementation by S.Ku., S.D.S., S.D. and K.L. Manuscript writing by S.Ku., S.D.S., A.C., S.D. and K.L. All authors reviewed the manuscript and approved the final version of the manuscript.

