## Supplementary Material for "Recognition determinants of improved HIV-1 neutralization by a heavy chain matured pediatric antibody"

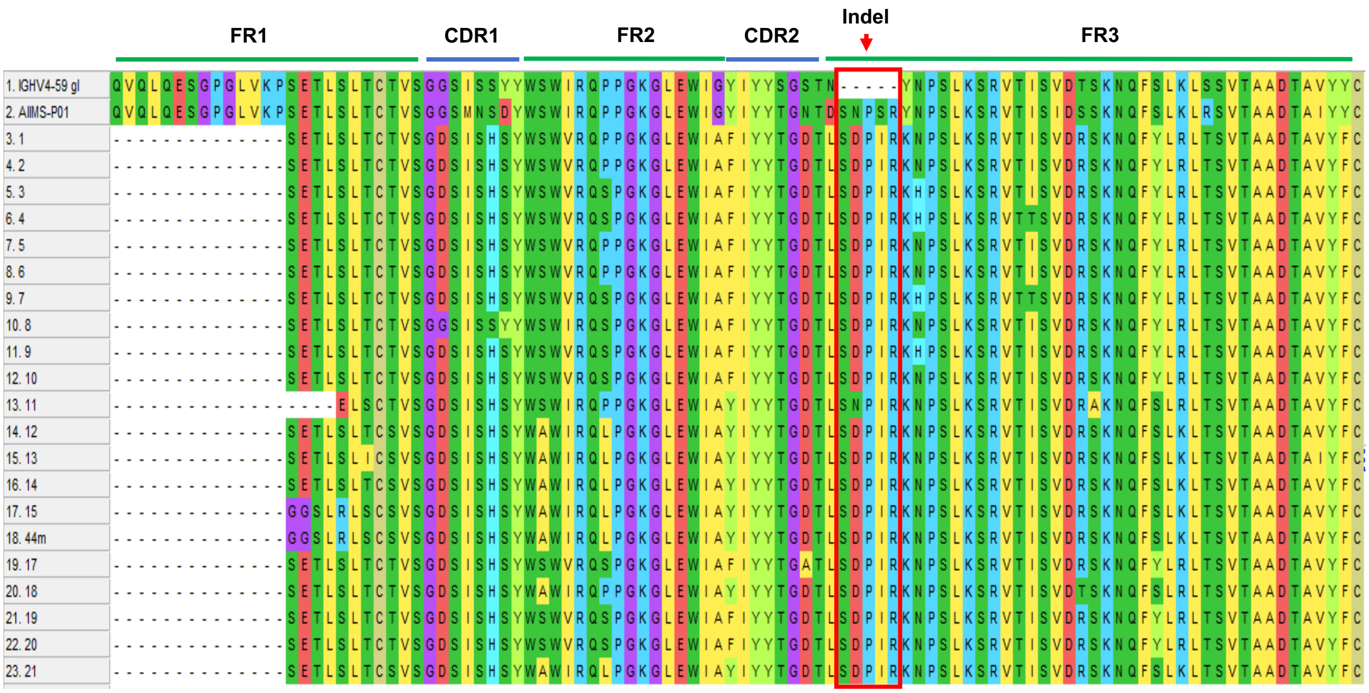

**Figure S1: Sequence analysis of HIV-1 pediatric bnAb AIIMS-P01 lineage members.** The alignment of 21 mutated members to the WT antibody AIIMS-P01, with highlighted indels. Antibody Framework Regions 1-3 (FRs) and Complementarity Determining Regions 1-2(CDRs) and indel have been marked. Related to Figure 1.

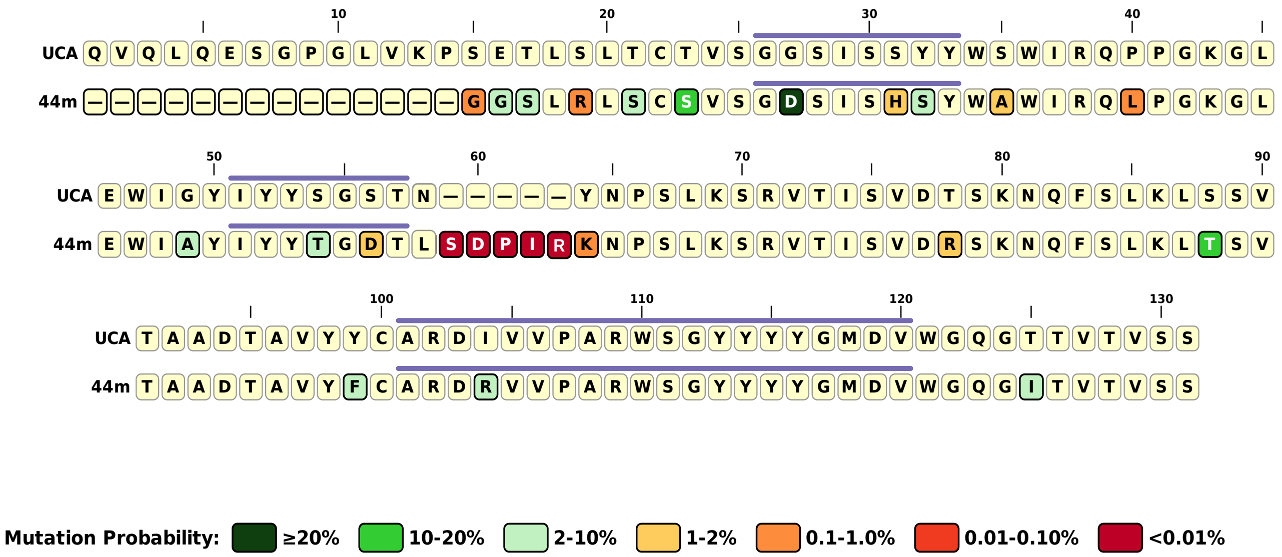

**Figure S2: Mutation analysis of pediatric HIV-1 bnAb 44m**. Related to Figure 1.

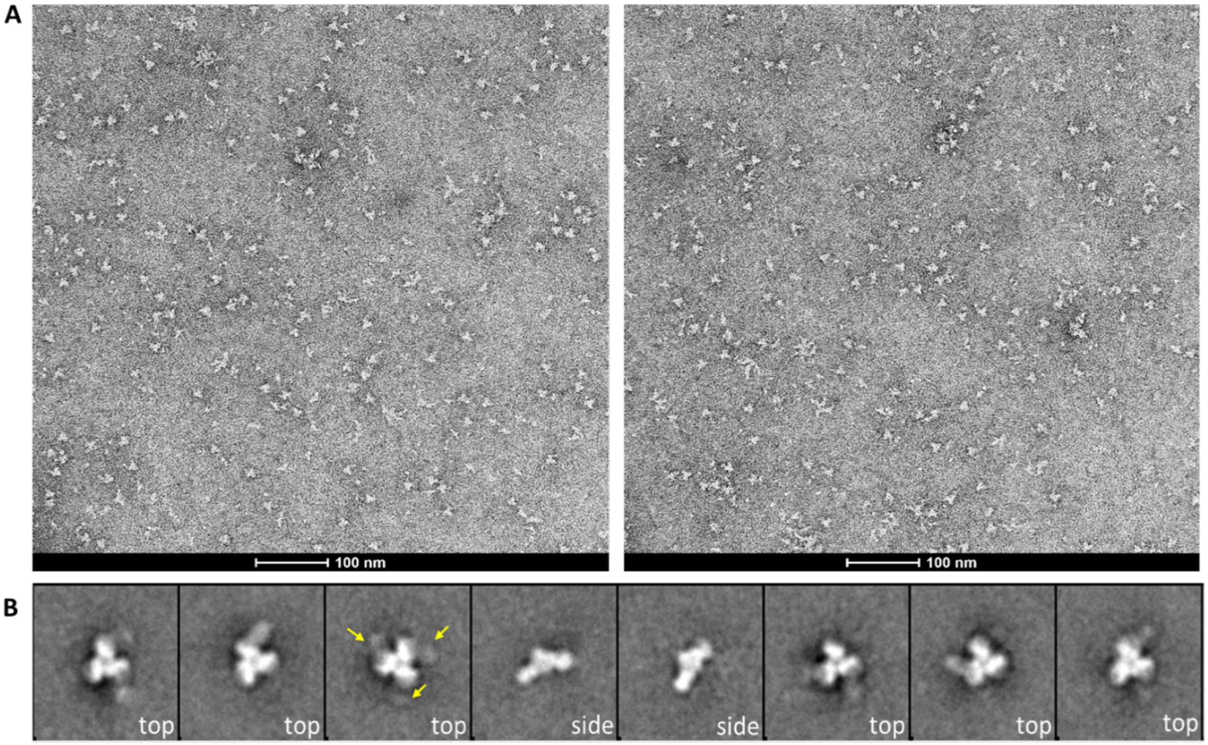

**Figure S3: Negative staining and 2D class averages of BG505.SOSIP.664.T332N gp140 Env trimer with 44m Fab. (A)** Representative negative-stain TEM (nsTEM) micrographs of BG505.SOSIP.664.T332N gp140 trimer and 44m Fab show homogeneous distribution of the SEC purified complex without any aggregation. **(B)** Representative reference-free 2D class averages of the nsTEM images indicate the emergence of extra densities at the apex of the trimeric gp120 protein. The extra densities corresponding to 44m bnAb are highlighted using yellow arrows. Related to Figure 3.

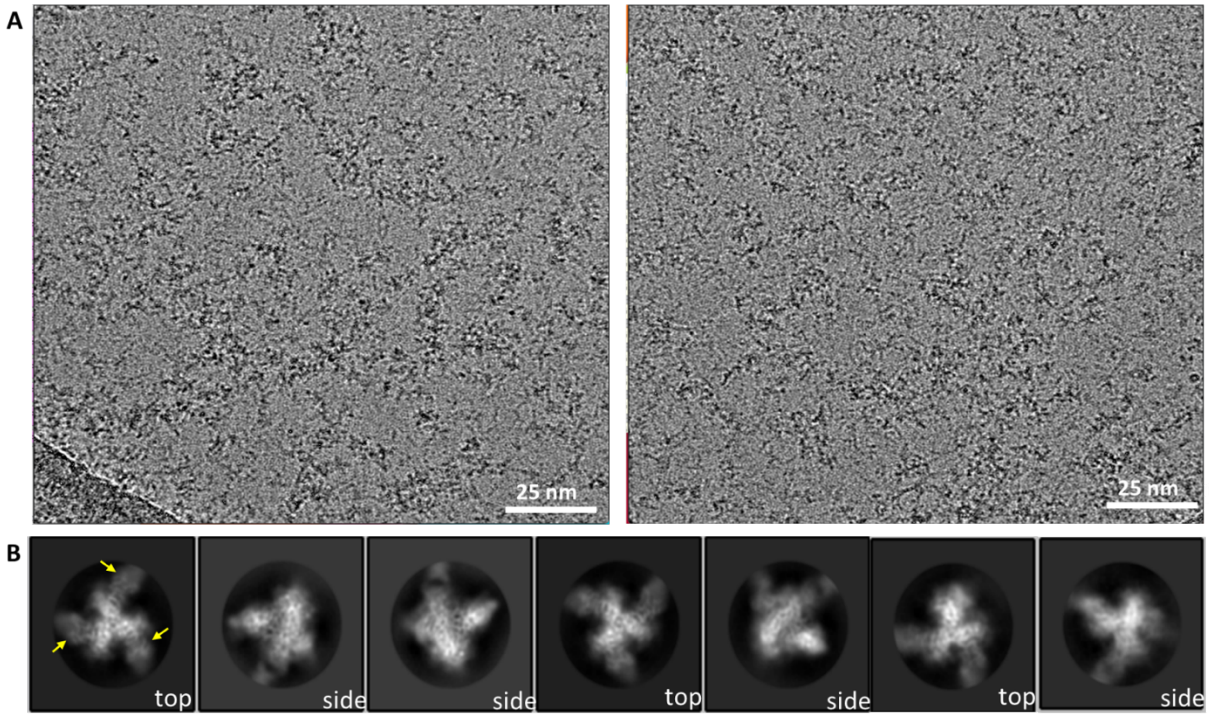

**Figure S4: Cryo-EM micrograph and 2D class averages of BG505.SOSIP.664.T332N gp140 Env trimer with 44m Fab. (A)** Representative Cryo-EM raw micrographs of BG505.SOSIP.664.T332N gp140 trimer and 44m bnAb ensure the particle homogeneity and equal distribution of the complex in near-native cryogenic temperature. **(B)** Representative reference-free 2D class averages of the nsTEM images clearly indicate the emergence of extra densities at the apex of the trimeric gp120 protein. Different views of the ternary complex having high-resolution features denote similar binding sites for 44m bnAb. The extra densities corresponding to 44m bnAb are highlighted using yellow arrows. Related to Figure 3.

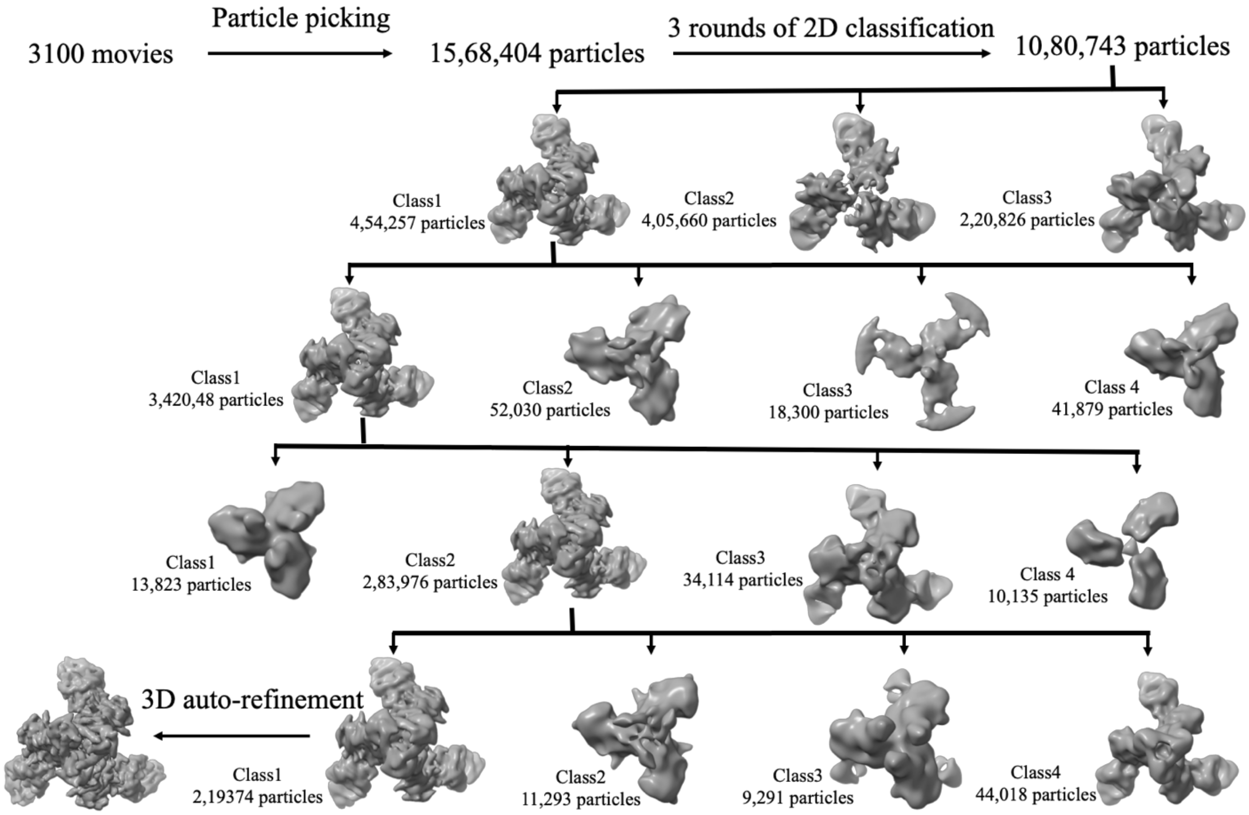

**Figure S5: Pipeline of data processing using single particle Cryo-EM and 3D classification BG505.SOSIP.664.T332N gp140 Env trimer in complex with 44m bnAb.** Cryo-EM data processing workflow and structure determination of BG505.SOSIP.664.T332N gp140 Env trimer in complex with 44m bnAb. Class 1 map after four rounds of 3D classification has high-resolution features, thus further refined and sharpened. Related to Figure 3.

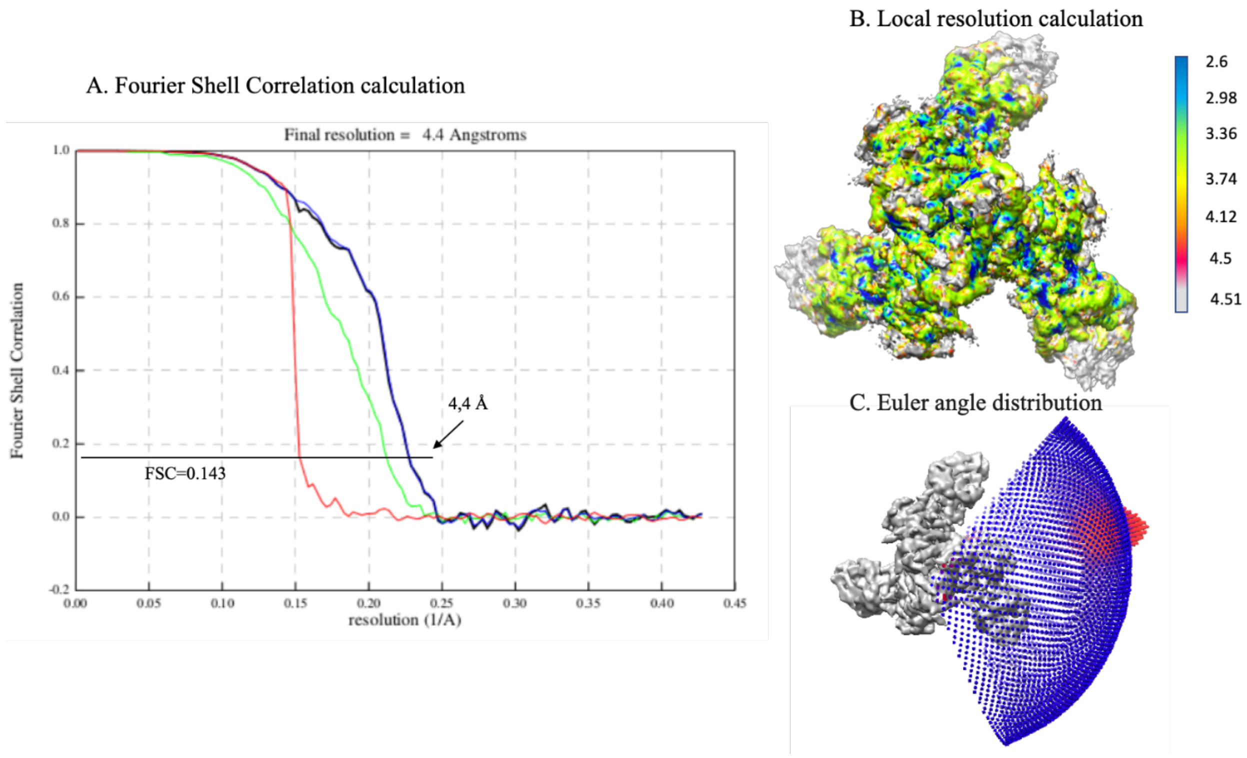

**Figure S6: High resolution 3D model reconstruction for BG505.SOSIP.664.T332N gp140 Env trimer in complex with 44m bnAb. (A)** Gold standard Fourier Shell Correlation (FSC) curve of the high resolution Cryo-EM map of BG505.SOSIP.664.T332N gp140 Env trimer in complex with 44m bnAb. **(B)** Local resolution calculation of Env trimer in complex with 44m bnAb using ResMap. **(C)** Euler angle distribution of particles contributing to form this trimeric complex. Related to Figure 3.

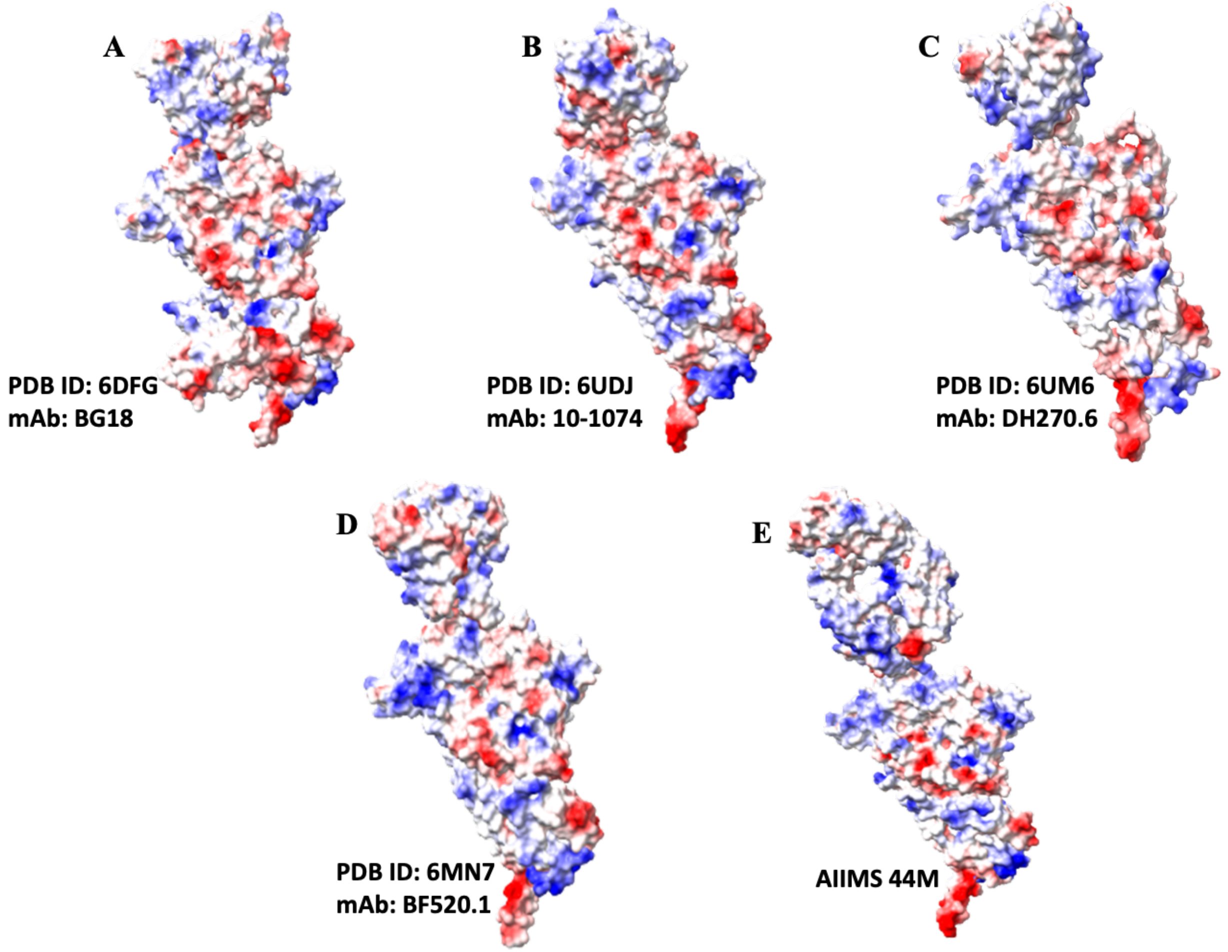

**Figure S7. Electrostatic surface potential map of different V3-glycan N332 targeting HIV-1 bnAbs in complex with Env trimers. (A-E)** The interacting surface indicates that the positioning of polar residues in the binding region facilitates the stabilization of the Env trimer 44m complex as reported in previous structures. Related to Figure 3 and 4.

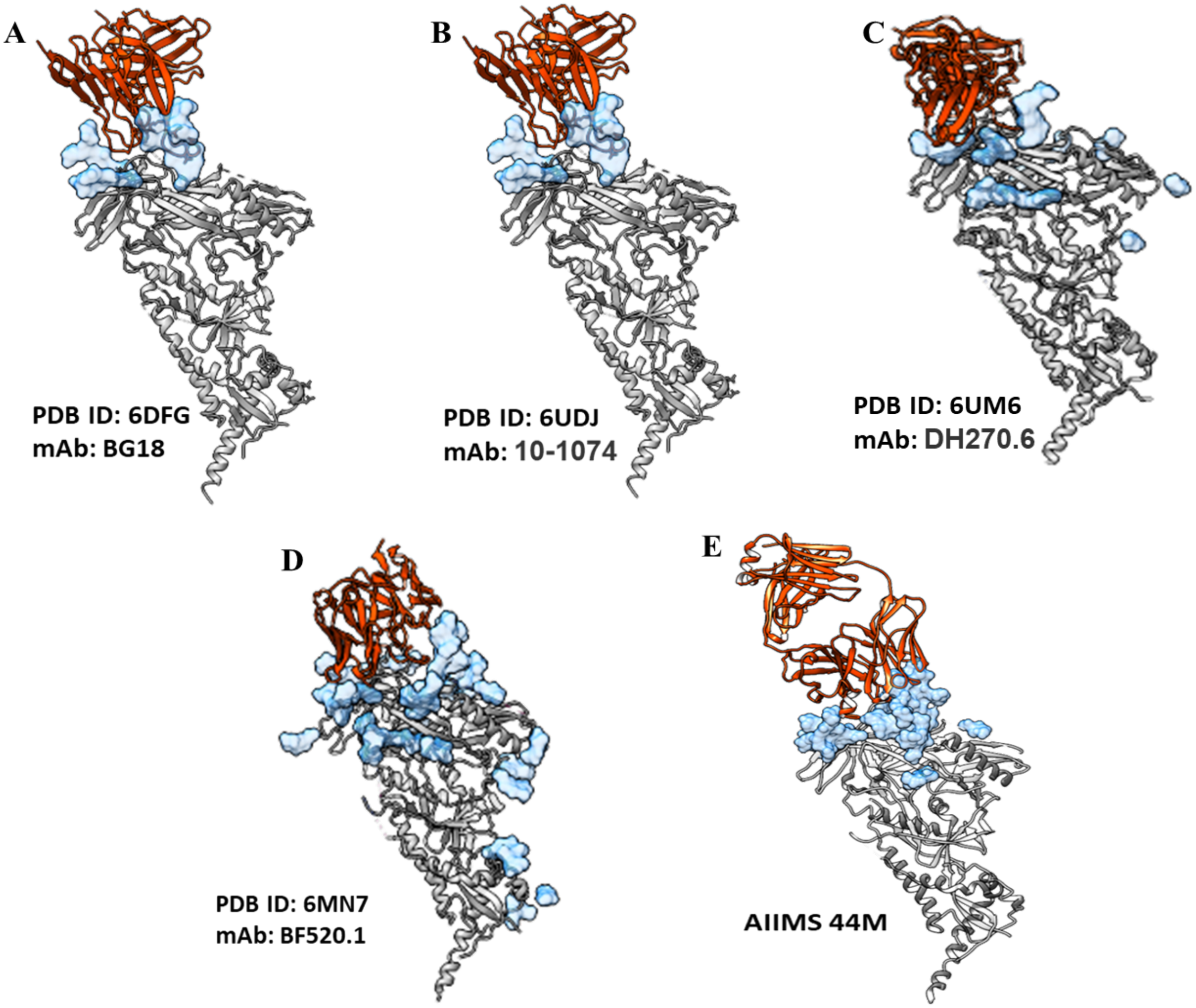

**Figure S8. Comparison of differential glycan binding pattern of various env trimer and V3-glycan N332 targeting HIV-1 bnAb complexes.** ***A-E.*** Different atomic models gp120 and bnAb showing different binding regions of the paratopes also unique glycan binding pattern. The glycan surface area in the antibody binding region is significantly higher in the case of 44m pediatric bnAb. Related to Figure 4 and 5.

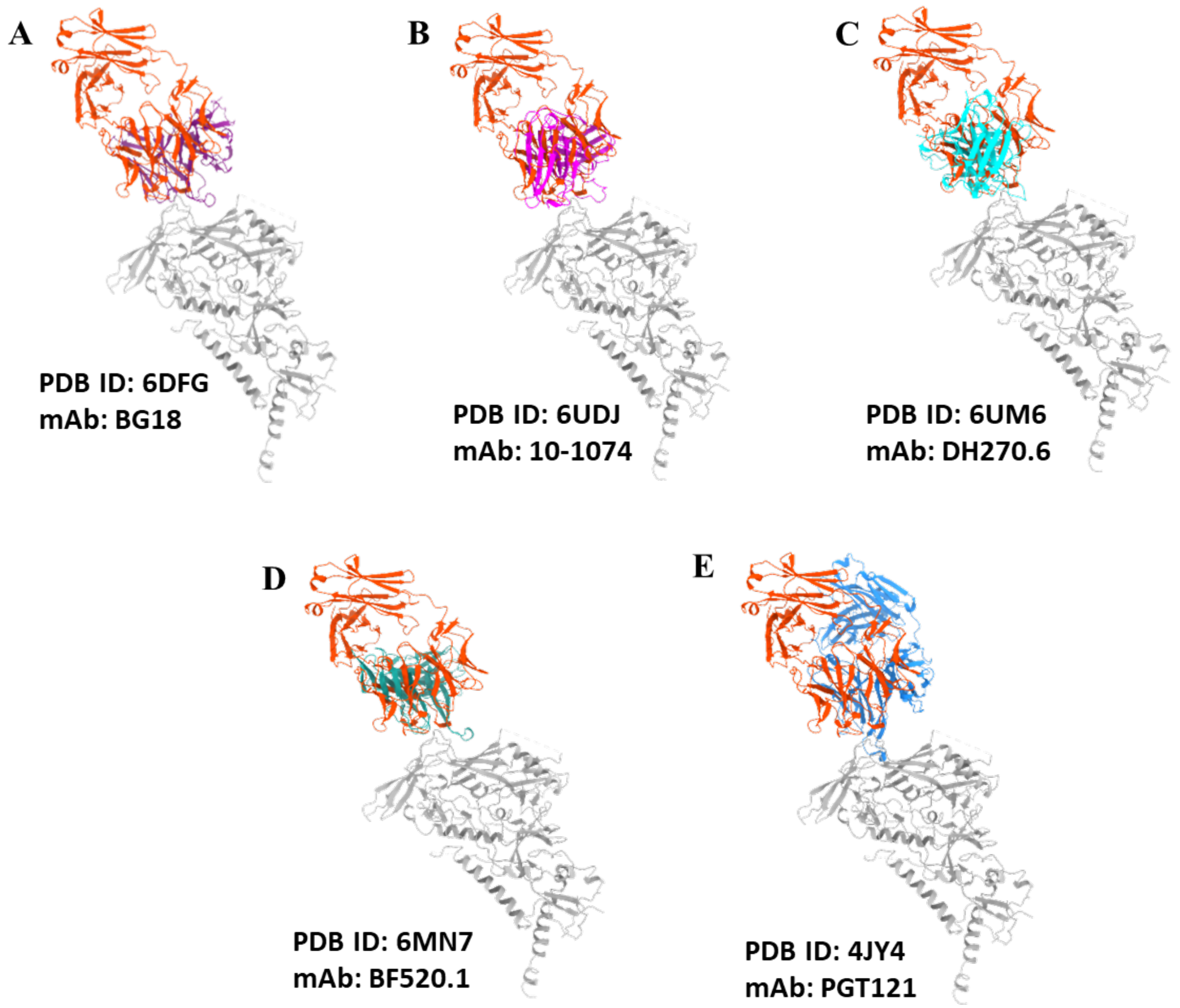

**Figure S9: Overlay of different V3-glycan N332 targeting HIV-1 bnAbs with 44m pediatric bnAb. A-E.** Superimposition of the antibodies binding in the V3 loop of the gp120 protomer shows that a common interaction pattern across all the bnAbs in the CD4 binding site. Related to Figure 4 and 5.

**A**

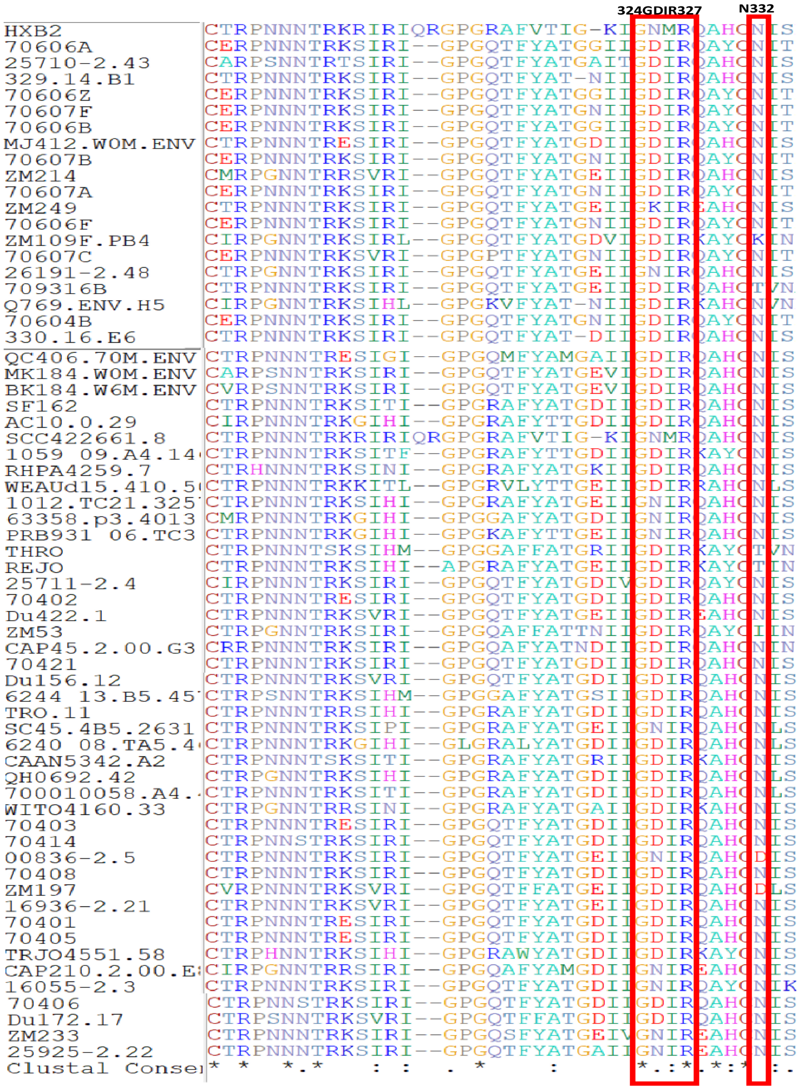

**B**

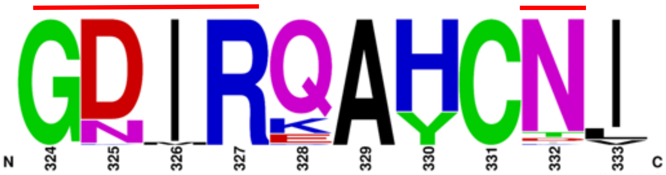

**Figure S10: Sequence analysis of HIV-1 viruses tested in neutralization assays. (A)** V3 loop sequence alignment **(B)** Logogram representing the frequency of the amino acid residues in V3 region GDIR motif and N332 highlighted in red line. Related to Figure 2, 4 and 5.

| | AIIMS P01 (WT) | AIIMS 44m (mutant) |
|---|---|---|
| **25710 WT** | 1.01 µg/ml | 0.45 µg/ml |
| **25710 D325N** | >10 µg/ml | 1.9 µg/ml |
| **Fold Decrease $IC_{50}$** | 44 | 4.2 |

| | | |
|---|---|---|
| **25710 D325K** | >10 µg/ml | >10 µg/ml |
| **Fold Decrease $IC_{50}$** | 56 | 59 |

**Figure S11: Neutralization $IC_{50}$ values of AIIMS-P01 WT and its lineage member 44m bnAb for their D325 dependence.** Related to Figure 2, 4 and 5.

**Table S1: Number of sequences in 2018 sample timepoint obtained from AIIMS_330 after filtering of deep sequencing data as described in Kumar S et.al., 2023.** Related to Figure 1.

| Group | Total number of sequences used for analysis |
|---|---|
| 330_2018 | 400249 |

**Table S2: Characteristics features of matured lineage members that share the same clonotype with AIIMS-P01.** Related to Figure 1.

| V-Gene | D-Gene | J-Gene | CDR3 length | CDR3 aa | Amino acid mutations | Isotype |
|---|---|---|---|---|---|---|
| IGHV4-59*01 | IGHD2-21*02 | IGHJ6*01 | 20 | ARDKVVPARWAGYYNYGMDV | 15 | IgG3 |
| IGHV4-59*01 | IGHD2-21*02 | IGHJ6*01 | 20 | ARDKVVPARWAGYYNYGMDV | 15 | IgG1 |
| IGHV4-59*01 | IGHD2-21*02 | IGHJ6*01 | 20 | ARDKVVPARWAGYYNYGMDV | 17 | IgG1 |
| IGHV4-59*01 | IGHD2-21*02 | IGHJ6*01 | 20 | ARDKVVPARWAGYYNYGMDV | 18 | IgG1 |
| IGHV4-59*01 | IGHD2-21*02 | IGHJ6*01 | 20 | ARDKVVPARWAGYYNYGMDV | 15 | IgG3 |
| IGHV4-59*01 | IGHD2-21*02 | IGHJ6*01 | 20 | ARDKVVPARWAGYYNYGMDV | 15 | IgG1 |
| IGHV4-59*01 | IGHD2-21*02 | IGHJ6*01 | 20 | ARDKVVPARWAGYYNYGMDV | 18 | IgG1 |
| IGHV4-59*01 | IGHD2-21*02 | IGHJ6*01 | 20 | ARDKVVPARWAGYYNYGMDV | 12 | IgG1 |
| IGHV4-59*01 | IGHD2-21*02 | IGHJ6*01 | 20 | ARDKVVPARWAGYYNYGMDV | 17 | IgG1 |
| IGHV4-59*01 | IGHD2-21*02 | IGHJ6*01 | 20 | ARDKVVPARWAGYYNYGMDV | 16 | IgG1 |
| IGHV4-59*01 | IGHD2-21*02 | IGHJ6*01 | 20 | ARDKVVPLQWSGYVYYGMDV | 14 | IgE |
| IGHV4-59*01 | IGHD6-25*01 | IGHJ6*01 | 20 | ARDRVVPARWSGYYYYGMDV | 14 | IgE |
| IGHV4-59*01 | IGHD6-25*01 | IGHJ6*01 | 20 | ARDRVVPARWSGYYYYGMDV | 16 | IgG1 |
| IGHV4-59*01 | IGHD6-25*01 | IGHJ6*01 | 20 | ARDRVVPARWSGYYYYGMDV | 14 | IgG1 |
| IGHV4-59*01 | IGHD6-25*01 | IGHJ6*01 | 20 | ARDRVVPARWSGYYYYGMDV | 19 | IgG3 |
| IGHV4-59*01 | IGHD6-25*01 | IGHJ6*01 | 20 | ARDRVVPARWSGYYYYGMDV | 19 | IgG1 |
| IGHV4-59*01 | IGHD2-21*02 | IGHJ6*01 | 20 | ARDKVVPARWAGYYNYGMDV | 16 | IgG1 |
| IGHV4-59*01 | IGHD2-15*01 | IGHJ6*01 | 20 | ARDRVVPAKWAGYYYFGMDV | 12 | IgG1 |
| IGHV4-59*01 | IGHD2-21*02 | IGHJ6*01 | 20 | ARDKVVPARWAGYYNYGMDV | 16 | IgG1 |
| IGHV4-59*01 | IGHD2-21*02 | IGHJ6*01 | 20 | ARDKVVPARWAGYYNYGMDV | 16 | IgG1 |
| IGHV4-59*01 | IGHD6-25*01 | IGHJ6*01 | 20 | ARDRVVPARWSGYYYYGMDV | 14 | IgG1 |

**Table S3: Cryo-EM data collection and processing.** Related to Figure 3.

| **Data collection and processing** | |
|---|---|
| Magnification | 42000 |
| Voltage (kV) | 200 |
| Electron exposure (e–/Å2) | 60 |
| Number of frames | 20 |
| Defocus range (μm) | -0.75 to –2.25 |
| Pixel size (Å) | 1.17 |
| Symmetry imposed | C3 |
| Number of movies | 3100 |
| Number of particles in model | 21 |
| Map resolution (Å) | 4.42 |
| FSC threshold | 0.143 |
| Map sharpening B factor (Å2) | -150 |
| **Validation** | |
| MolProbity Score | 2.12 |
| Clash Score | 14.11 |
| Rotamer Outliers (%) | 0 |
| Ramachandran Plot | |
| Outlier (%) | 0 |
| Allowed (%) | 7.31 |
| Favored (%) | 92.69 |
| EMRinger Score | 1.90 |
